# Neutralization activity in chronic HIV infection is characterized by a distinct programming of follicular helper CD4 T cells

**DOI:** 10.1101/2024.07.31.605954

**Authors:** Eirini Moysi, Ashish A. Sharma, Sijy O’Dell, Spiros Georgakis, Perla Mariana Del Rio Estrada, Fernanda Torres-Ruiz, Mauricio González Navarro, Yara Andrea Luna Villalobos, Santiago Avila Rios, Gustavo Reyes-Teran, Margaret H. Beddall, Sung-Hee Ko, Frida Belinky, Michail Orfanakis, Laurence de Leval, Ana B. Enriquez, Clarisa M. Buckner, Susan Moir, Nicole Doria-Rose, Eli Boritz, John R. Mascola, Rafick-Pierre Sekaly, Richard A. Koup, Constantinos Petrovas

## Abstract

A subset of people living with HIV (PLWH) can produce broadly neutralizing antibodies (bNAbs) against HIV, but the lymph node (LN) dynamics that promote the generation of these antibodies are poorly understood. Here, we explored LN-associated histological, immunological, and virological mechanisms of bNAb generation in a cohort of anti-retroviral therapy (ART)-naïve PLWH. We found that participants who produce bNAbs, termed neutralizers, have a superior LN-associated B cell follicle architecture compared with PLWH who do not. The latter was associated with a significantly higher *in situ* prevalence of Bcl-6^hi^ follicular helper CD4 T cells (TFH), expressing a molecular program that favors their differentiation and stemness, and significantly reduced IL-10 follicular suppressor CD4 T cells. Furthermore, our data reveal possible molecular targets mediating TFH-B cell interactions in neutralizers. Together, we identify cellular and molecular mechanisms that contribute to the development of bNAbs in PLWH.

## Introduction

Chronic HIV infection changes the microarchitecture and cellular composition of LNs (*1*). Microanatomical transformations include disruptions to follicular reticular cell conduits (*2*), and dendritic cell (FDC) networks(*3*), progressive loss of follicular B-cell architecture (*4*), and alterations in the phenotype and function of T and B cells that can persist despite antiretroviral therapy (ART)(*5–8*). Follicular helper T cells (TFH) cells are a subset of CD4 T cells defined by low expression of CCR7 and high expression of the chemokine receptor CXCR5, costimulatory receptors PD-1 and ICOS(*9*), and lineage commitment transcription factor Bcl-6(*10*). Among other functions, these cells are critical for the initiation and maintenance of germinal center reactions which is where somatic hypermutation – a process necessary for the generation of high-affinity antibodies-takes place(*11*). Non-human primate (NHP) studies of SIV infection have found that TFH cells can be detected as early as 14 days post-challenge(*12*). Furthermore, CXCR5^hi^PD-1^hi^ CD4+ T cells accumulate in the blood and LNs in the chronic stage of the disease in NHPs (*13*) as well as in humans (*14*). Despite this enrichment, disease-associated impairments in TFH cell function have been reported, and people living with HIV (PLWH) have diminished protective antibody responses as (*5, 15*). However, whether such functional impairments correlate with altered B cell follicle architectures and to what extend the latter affect the development of cross-neutralizing and broadly neutralizing antibodies (bNAbs) against the virus itself in chronic HIV disease remains unclear.

Generation of bNAbs is a critical goal of HIV vaccination strategies as these can neutralize diverse viral isolates. These antibodies have been found to arise in a subset of PLWH (10-30%) following 2-3 years of infection (*16*), exhibit unusual structural changes to the antigen binding regions and surrounding framework regions and have a high degree of somatic hypermutation(*17–19*). High HIV plasma viremia, low total CD4 T cell counts, high frequencies of TFH cells early in infection and low frequencies of regulatory T cells (T_REG_) have all been associated with induction of bNAbs in PLWH (*20–23*). Extrafollicular (T_REG_) and follicular (T_FR_) regulatory CD4 T cells accumulate in the lymph nodes of untreated PLWH and suppress TFH and B cells, which impairs germinal center reactivity and IgG production (*24*). How these dynamics translate at tissue level in the context of chronic HIV viremia in individuals with and without broad neutralizing antibody production, however, remains poorly understood.

In this study, we sought to address these questions by characterizing TFH, T_REG_ cells and overall GC structure in PLWH. We performed a detailed characterization of LN-architecture and used flow cytometry, multiparametric confocal imaging and single-cell RNA sequencing to elucidate the virological and immunologic signatures associated with the production of cross-neutralizing antibodies. Our results reveal that PLWH with high antibody neutralizing activity, termed “neutralizers”, have higher frequencies of T central memory cells and TFH cells and unique B cell follicle architecture compared to non-neutralizers. We observed that active, polarized GCs were more frequent in the tissues of neutralizers and contained transcriptionally distinct TFH that could account for the neutralization activity between the two groups of PLWH. Together, our results elucidate mechanisms which contribute to the generation of neutralizing antibodies in PLWH. These insights can be used to develop therapeutics and vaccination strategies that promote the generation of bNAbs in PLWH.

## Results

### Determination of plasma neutralizing activity in study participants

To determine whether differences exist between profiles with neutralizing and non-neutralizing antibodies, we performed an analysis of cross-neutralizing activity in the serum of a cohort of 147 ART-naïve people living with chronic HIV infection. Sera were initially screened against a mini-panel that consisted of the following six isolates: Q259.17 (A), Q461.e2 (AD), TH976.17 (AE), YU2.DG (B), CH070.1 (BC) and 0013095-2.11 (C), and findings were then validated for a subset of sera (n = 62) with an extended panel of 20 HIV-1 isolates from clades A, AD, AE, AG, B, BC, C and D. A range of cross-neutralizing activities was observed in the mini-panel screenings. As shown in **Fig. 1A**, cross-neutralization of 50% of isolates (3 or more) was observed in 23% of sera, whilst 76% of sera had either low (16-33% of isolates) or no measurable activity. A similar pattern was observed in the extended panel screenings **(Fig. S1A).** Cross-neutralization of 45% of isolates or more at an ID_50_>40 was observed in 25% of sera (high activity), whilst 20% of sera could cross-neutralize 20-40% of isolates (medium activity) and 50% of sera <15% of isolates (low or no activity).

**Fig. 1:**
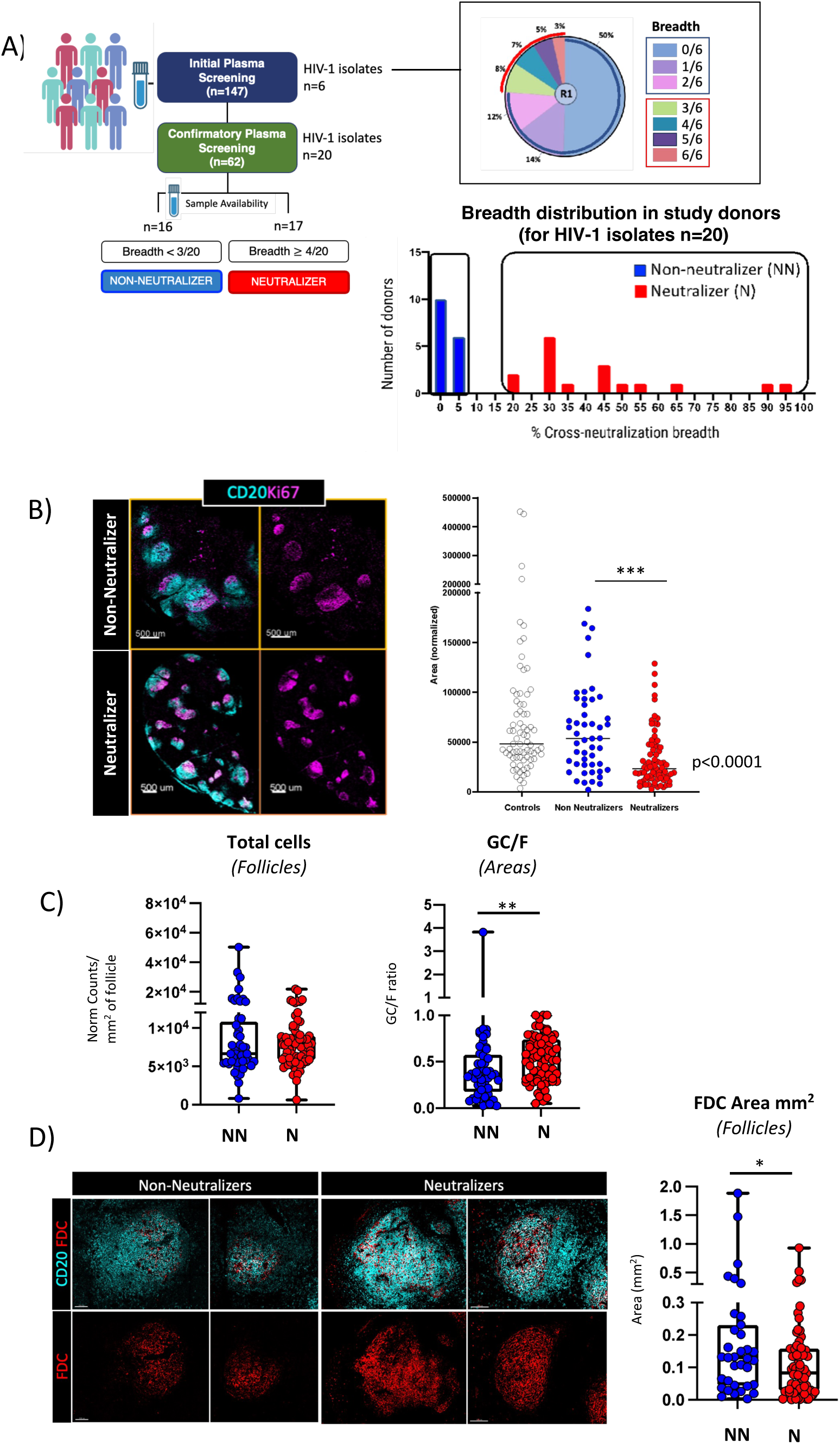
Cross-neutralization is associated with GCs displaying preserved features of maturation. A) Diagram depicting the strategy employed for neutralization screening (left) as well as the distribution of cross-neutralization breadth in the population of study, defined as the percentage of the number of isolates recognized with an ID_50_>40 out of 6 isolates (pie chart) and 20 isolates (bar graph) tested among study participants. B) Representative confocal images showing the distribution of CD20 (cyan) and Ki67 (magenta) in NNs and Ns and total B-cell follicle area quantification (CD20^hi/dim^) in control tissues, NNs and Ns. Each circle represents a follicle, and data are for all tissues combined (***p<0.001). Original magnification x40 (NA 1.3), scale 500um. C) Box plots showing the number of total cells and GC/F ratios (areas) in NN and Ns as measured by quantitative imaging analysis and Histocytometry. Data for each follicle represent total counts of JOJO-1+ events (total cells) normalized to the total area (mm^2^) of follicle screened. Each circle represents an individual follicle, and data shown are for all follicles combined. p=0.4249, **p=0.00115. D) Representative confocal images showing the distribution of FDC (red) in B cell follicles (CD20^dim/hi^ areas, cyan) in NNs vs Ns (original magnification x40, scale 100um), and box plot summarizing the FDC areas (mm^2^) in NNs and Ns (pooled data) as calculated using data from quantitative imaging analysis and Histocytometry (*p=0.0352). Each circle represents and individual follicle, and data shown are for all follicles combined. p values were considered significant if p≤0.05.

Based on these findings, and overall tissue and lymph node cell suspension sample availability, a total of 33 participants without (n = 16) or with (n = 17) cross-neutralizing activity were selected for further study (**Table 1, Table S1 and Table S2**). Participants were categorized as neutralizers (Ns) if their serum neutralizing activity in the extended panel was medium or high (>20% of isolates), or non-neutralizers (NNs) if their serum neutralization activity was less than 5% (**Fig. 1A, Fig. S1A and Fig. S1B**). Since neutralization breadth has been previously shown to depend on the nature of bNAb lineages involved (*25*), we sought to test whether known bNAb lineages were represented in the Ns. We performed computational fingerprinting analysis using a neutralization fingerprinting algorithm (NFP) that predicts the prevalence of each of ten reference epitope-specific bNAb groups(*26*). A total of five bNAb groups, VRC01-, 8ANC195-, PG128-, 10E8- and PGT151-like, were detected with a prevalence coefficient value ≥ 0.25 which is typically considered positive **(Fig. S1C)**. Therefore, serum neutralizing activity in Ns was consistent with the presence of known bNAb lineages.

**Table 1:** Clinical characteristics of study participants.

| Donor group | No. of donors | Median (range) |  |  |  |
| --- | --- | --- | --- | --- | --- |
|  |  | Age | LogVL | CD4+ cells/ul | % CD4 |
| **ALL** | 33 | 29 (18-61) | 5.13 (1.6-7) | 348 (4-716) | 17 (1-37) |
| **Non-neutralizers** | 16 | 26 (18-40) | 5.42 (1.6-7) | 415 (153-716) | 24 (9-37) |
| **Neutralizers** | 17 | 31 (20-61) | 4.96 (3.65-5.76) | 279 (4-671) | 12 (1-22) |

### Cross-neutralization is associated with a greater degree of B-cell follicle preservation and GC formation

HIV acquisition causes morphological changes in the microarchitecture of LNs that affect the ability of B cell follicles to perform their function over time (*27*). To understand if such anatomical changes could be associated with reduced cross-neutralization in our study, we examined the features of B-cell follicle anatomy that were differentially represented in participants with and without serum cross-neutralizing activity. First, the area of follicular Regions of Interest (ROIs), defined by the expression of CD20 (**Fig. 1B** and **Fig. S1D**) was evaluated in our cohort and compared to benign LN from HIV-negative participants that were characterized by mature follicles (follicular hyperplasia) (**Table S1**). The calculated areas were found significantly higher in the control and non-neutralizing compared to neutralizing participants, indicating a higher degree of follicular hyperplasia in non-neutralizers, in line with previous data (*4*) (**Fig.1B** and **Fig. S1D**). Although not significant, a trend for higher circularity of follicular ROIs was found in N compared to NN participants (**Fig. S1D**).

We then evaluated additional elements associated with HIV disease including the overall B cell follicle cellularity, extend of Ki67^hi/lo^ CD20+ areas as a surrogate of GCs, and FDC network preservation(*28, 29*). We found a similar degree of overall cellularity in Ns–defined as the total number of nucleus positive cells measured within CD20^hi/dim^ B cell follicles–as compared with NNs **(Fig. 1C).** Next, we examined the extent of GC preservation amongst tissues. For the quantitation, we focused on Ki67, a marker of proliferation and cell cycle progression(*30*), as lack of Ki67 polarization in HIV GCs has previously been shown to associate with GC disorganization(*31*). Analysis of the areas occupied by CD20^hi/dim^ Ki67^hi^ B-cells (denoted as GCs) versus CD20^hi/dim^ Ki67^hi/lo^ B cell areas (denoted here as total follicle or F) revealed a significantly higher ratio of GC/F in Ns compared with NNs (p=0.00115) **(Fig. 1C** and **Fig. S1E).** Next, we extended our analysis to FDC networks and observed that despite the significantly higher FDC areas in NNs as compared with Ns (p=0.0352) (**Fig.1D**), a reversed trend was found for the FDC/F ratios, but the difference was not statistically significant (**Fig. S1F and Fig. S1G**). Together, these findings suggest that neutralization activity in chronic HIV is associated with better preserved follicular structures.

### Cross-neutralization is associated with distinct phenotypic signatures of TFH cells

Having observed morphological differences in the LNs of non-neutralizers and neutralizers, we next wanted to assess the cellular composition of the LNs in these two groups. We performed flow cytometry on single cell suspensions of LNs and enumerated the frequencies of the following immune subsets: naïve CD4 T cells [CD27^lo^CD45RO^lo^], total memory [CD27^hi/lo^CD45RO^hi^], central memory [T_CM_; CD27^hi^CD45RO^hi^], effector memory [T_EM_; CD27^lo^CD45RO^hi^] and TFH cells (PD-1^hi^CXCR5^hi^) (**Fig. 2A and Fig. S2A)**. We found a significantly higher percentage of total memory T cells in neutralizer LNs compared to non-neutralizers (p=0.032) **(Fig. 2B)**. This was primarily driven by an increase in T_CM_ cells (p = 0.005) **(Fig. 2B)**. Furthermore, we observed a positive association between the latter frequencies and the recorded neutralization potencies which although not significant (R^2^=0.1654, p=0.1052) was suggestive of correlated changes between these two factors (**Fig. 2B**).

**Fig. 2:**
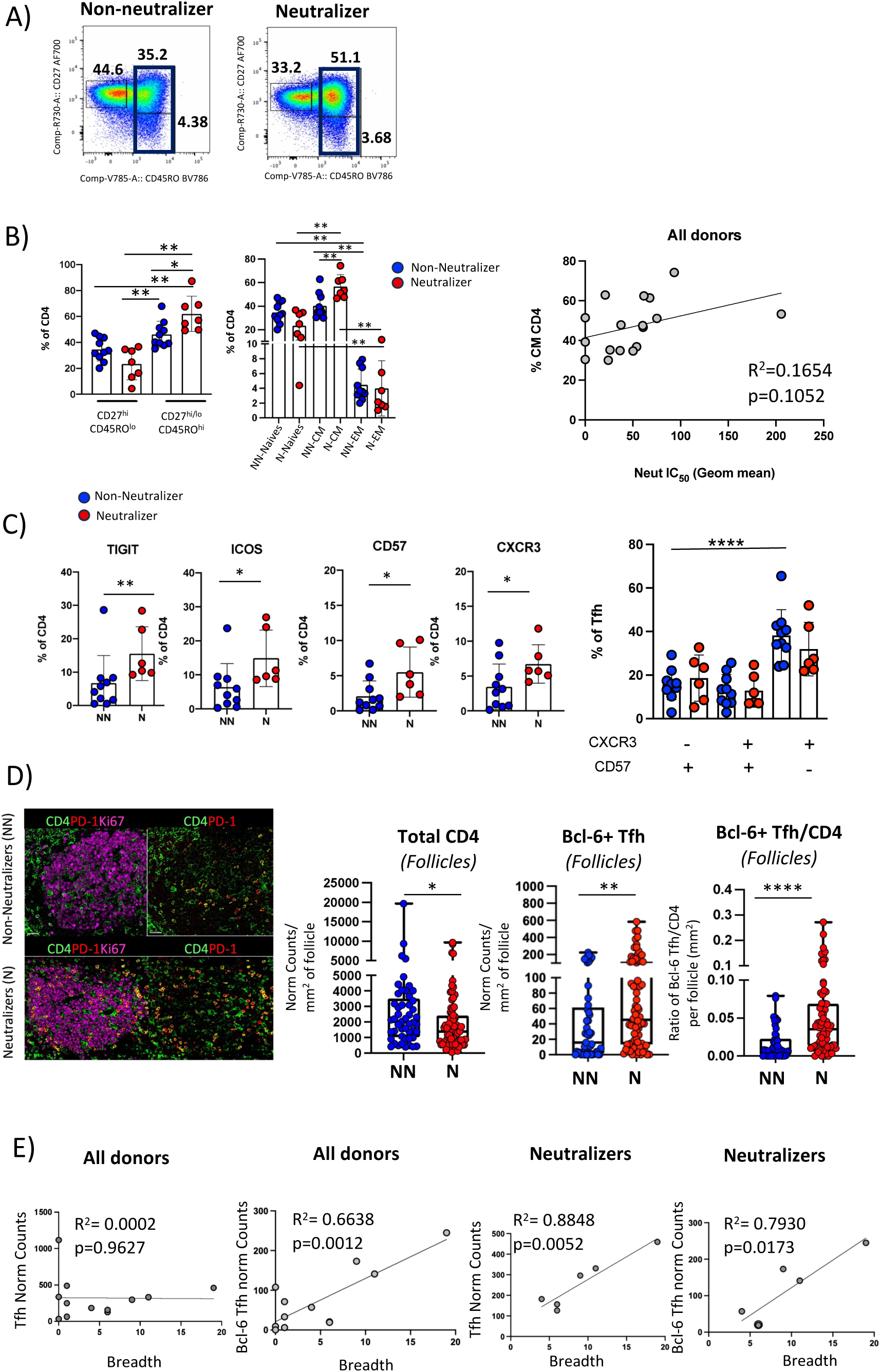
Cross-neutralization is associated with an expansion of the LN CD4 T cell memory compartment and a distinct TFH cell phenotypic profiling. A) Flow cytometry plots depicting the frequencies of memory CD4 T cell subsets in NNs and Ns within the CD3^hi^CD4^hi^ gated populations. B) Bar graphs showing the relative frequencies of CD4 T cell subsets in Ns and NNs (left panel) and the positive association between the % of T_CM_ CD4 T cells and micro-neutralization ID_50_ titers as assayed against a 20 HIV isolate panel. C) Cumulative flow cytometry plots showing the frequencies of receptors TIGIT, ICOS as well as CD57 epitope and CXCR3 within the PD-1^hi^ CXCR5^hi^ TFH cell compartment in NNs and Ns, and frequencies of combined CXCR3 CD57 expression. TIGIT **p=0.0075; ICOS, *p=0.0160; CD57 *p=0.0225, CXCR3 * p=0.0420 (Mann Whitney U test); NN CXCR5^lo^CD57^hi^ vs NN CXCR3^hi^ CD57^lo^ *p<0.0001 (ANOVA). D) Representative confocal images of GCs in NNs and N showing the distribution of TFH (CD4: green, PD-1^hi^: red) within the Ki67 follicular areas (magenta) and pooled histocytometry quantitation of total GC-CD4, GC Bcl-6^hi^ Tfh and GC Bcl-6^hi^ Tfh/CD4 ratios in NNs (n=6, blue circles represent all follicles counted in those six individuals) vs Ns (n=6, red circles represent all follicles counted as for NNs). The numbers of cells per mm^2^ of follicular area are shown. p values are *p= 0.0198 (ANOVA), **p= 0.003 and ****p<0.0001 (Mann-Whitney U test). Original magnification x40, scale bar = 50um. E) Regression plots showing the association between bulk TFH and Bcl-6^hi^ TFH cells (normalized cell counts adjusted for mm^2^ of area screened) as quantified by confocal imaging analysis and histocytometry, and the breadth of neutralization expressed as the number of HIV isolates recognized out of the 20 isolates tested.

We also found a trend for decreased non-TFH (PD-1^lo^CXCR5^lo^, PD1^lo^CXCR5^dim^) associated with increased pre-TFH (PD-1^dim^ CXCR5^lo^) and TFH cell frequencies in neutralizers compared to non-neutralizers (**Fig. S2B**). We then used flow cytometry to assess whether TFH related receptors were differentially expressed between neutralizers and non-neutralizers. Co-stimulatory receptors, such as CD226, ICOS, OX40 and co-inhibitory receptors, such as TIGIT, contribute to distinct functions of TFH cells(*32–34*). We also included in our analysis CD57, a glycan carbohydrate epitope expressed by a subset of TFH cells in the GC(*35, 36*) and CXCR3, a chemokine associated with a Th1-like TFH cell signature. Our analysis revealed a significant increase in the frequency of TFH cells expressing TIGIT (p = 0.0075), ICOS (p = 0.0160), CD57 (p = 0.0225) and CXCR3 (p = 0.0420) in Ns compared with NNs **(Fig. 2C).** Increases in CD226 (p = 0.0075) and CD95 (p = 0.0312) within the CD4 T cell compartment in Ns compared with NNs were also observed suggesting, along with TIGIT and ICOS a higher level of differentiation and activation in neutralizers **(Fig. S2C)**. TIGIT expression significantly correlated with the expression of ICOS and CD226 in both participant categories (R^2^=0.97, p<0.0001 for ICOS, and R^2^=0.62, p = 0.0003 for CD226) **(Fig. S2C)**. No significant differences, however, were observed with respect to the expression of PD-L1, CTLA-4, TIM-3, OX40 and OX40L between the two groups **(Fig. S2C).** Further probing of TFH heterogeneity revealed significantly higher frequencies of CD57^lo^ CXCR3^hi^ TFH cells compared to CD57^hi^ CXCR3^lo^ TFH cells in NNs (p<0.0001). A trend for higher frequencies of CD57^lo^ CXCR3^hi^ TFH cells in NNs as compared with Ns was also notable **(Fig. 2C)**. Similar findings were also observed when data from samples with sufficiently representative TFH cell numbers from NNs and Ns were dissected by t-Distributed Stochastic Neighbor Embedding (t-SNE) analysis **(Fig. S2D).** A total of 8 meta-clusters were identified by FlowSOM analysis displaying varying expressions of activating and inhibitory receptors **(Fig. S2D).** Whilst differences amongst the clusters did not reach statistical significance, there was a trend for higher representation of CD57^lo^CXCR3^hi^ TFH cells within the TIGIT^hi^ICOS^hi^CD95^hi^ phenotypic compartment in NNs compared with Ns (cluster/population 5) **(Fig. S2D)**. On the other hand, a trend for a higher frequency of the cluster bearing the CD57^hi^CXCR3^lo^ signature was seen in Ns (cluster/population 6) **(Fig. S2D).** Taken together, our data suggest an altered differentiation profile of TFH cells in non-neutralizers which is characterized by an accumulation of less differentiated (CD57^lo^), Th1-like (CXCR3^hi^) phenotype in NNs compared to Ns.

As location of TFH cells is necessary to understand function, we next analyzed the prevalence of TFH cells within CD20^hi/dim^ follicular areas of the LN *in situ*. Using immunofluorescence, we found significantly higher normalized numbers of bulk follicular CD4 T cells in NNs than Ns (p=0.0198) (**Fig. 2D and Fig. S3A)**. In contrast to total CD4 T cells, PD-1^hi^Bcl-6^hi^ TFH cells were significantly increased in Ns compared with NNs (p=0.0030) **(Fig. 2D and Fig. S3B).** Furthermore, the ratio of TFH to CD4 T cells within the B cell follicles of Ns was significantly higher compared to that of NNs (p<0.0001) **(Fig. 2D and Fig. S3B).** We also examined if a correlation existed between the normalized TFH cell counts and the breadth of serum cross-neutralization. Contrary to PD1^hi^ TFH cells, a significant correlation between PD-1^hi^Bcl-6^hi^ TFH cells and the cross-neutralization breadth in all participants (R^2^=0.66, p = 0.0012) was found (**Fig. 2E**). A similar profile was found when the group of neutralizers was analyzed separately (R^2^=0.79, p = 0.0173), which extended also to PD-1^hi^ TFH cells (R^2^=0.88, p = 0.0052) **(Fig. 2E).** Overall, these results show that the development of cross-neutralizing breadth is associated with a higher *in situ* prevalence of PD-1^hi^ TFH cells and ‘effector’ Bcl-6^hi^ TFH cells.

### Cross-neutralization is associated with a molecular profile favoring the development and function of TFH cells

To examine whether the transcriptomic profile of TFH cells is different between N and NN groups, we performed 10X single cell RNAseq (scRNAseq) on LN cells. Analysis of lineage biomarkers showed a significant expansion of B cells in Ns while a significantly higher frequency of bulk CD4 T cell was observed in NNs (**Fig. 3A** and **Fig. S4A**). Further clustering, based on the expression of CD4 T cell specific genes (**Fig. S4B**), revealed a high heterogeneity of the CD4 T cell compartment in both groups (**Fig. S4B**). We observed an increase of TFH cells in Ns and a significantly higher frequency of naïve CD4 T cells in NNs than Ns (**Fig. 3B**), which corroborates our flow cytometry data (**Fig. 2**). We then focused on characterizing the TFH cell compartment. We observed a higher transcription of genes encoding for critical regulators that promote TFH cell differentiation (e.g., c-Maf, IL6ST, Tox2), stemness (Tcf7) and interaction with GC B cells (CXCL13) (**Fig. 3C**) in NNs compared to Ns. Interestingly, TFH cells from Ns had a significant upregulation in IL-6 and Wnt signaling pathways as well as of targets which are downstream of Tcf7, SMAD3,4 and STAT3 while NN TFH cells are associated with significant upregulation of type I and II IFN signaling and STAT6 target genes (**Fig. 3C**). With respect to the regulatory CD4 T cell (T_REG_) transcriptomic profile, genes like FOXP1, a co-factor for FoxP3(*37*) and IRF1, an inducer of type 1 regulatory Tr1 cells (*38*) and negative regulator of TFH cell differentiation(*39*) were upregulated in NN compared to N participants (**Fig. 3D**). Furthermore, downstream targets for several established T_REG_ inducers including STAT3, 5, 6(*40–43*), c-Myc(*44*), SMAD3(*45*), Vitamin D Receptor(*46*) and FOXP3/FOXP1 were upregulated in NN compared to Ns (**Fig. 3D**). Our data indicate that despite the higher frequency of FOXP3^hi^ CD4 T cells in N, the NNs are characterized by a molecular profile favoring the development of potential suppressor CD4 T cells. When we validated molecules revealed by sc-RNA analysis with flow cytometry, we also observed significantly higher levels of combined TCF1^hi^AHR^hi^CAV1^hi^CD130^hi^ expression in TFH cell subsets (PD-1^hi^ CD57^lo/hi^), compared to naïve CD4 T cells (4.04±3.65% vs 0.21±0.13%, p = 0.0261) and compared to the PD-1^lo^CD57^lo^ CD4 T cell subset (4.04±3.65% vs 0.17±0.17%, p = 0.0244) in Ns compared to NNs **(Fig. S4C)**. In sum, our data suggests that TFH cells from Ns express a molecular program that favors their differentiation and stemness.

**Fig. 3:**
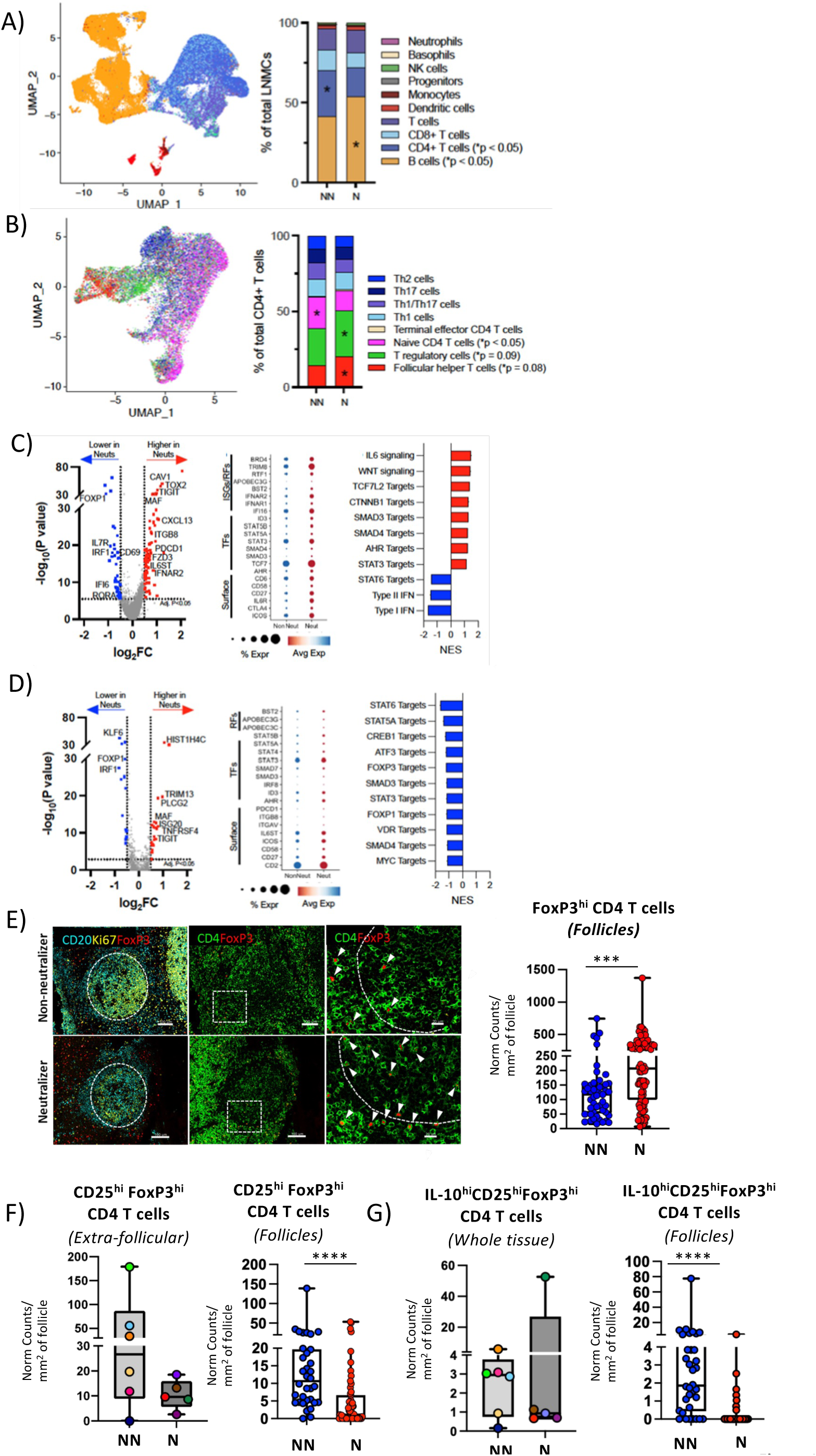
TFH cells are characterized by a molecular profile favoring their differentiation and longevity in cross-neutralization participants. UMAP projections of total LNMCs (A) and CD4 T cells (B), with color-coded cell populations and CD4 T cell subpopulations identified and their percentages in NNs vs Ns. C-D) Volcano plots and dot plots of differentially expressed genes (DEGs) and normalized enrichment scores (NES) of GSEA for selected signaling and downstream targets are shown. The DEGs were calculated using the MAST (Hurdle model based) test via the FindMarkers function using the Seurat package in R. The p-values were corrected using the Benjamini-Hochberg method and genes meeting an adjusted p< 0.05 and abs(log_2_FC)>0.5 are highlighted on the volcano plot. Blue dots and red dots in volcano plots represent genes that are significantly downregulated or upregulated in neutralizers respectively. E) Representative confocal images showing FOXP3^hi^ CD4 T cells in extrafollicular and follicular areas in NNs vs N. FOXP3^hi^ CD4 T cells (white arrows) were defined by means of concurrent CD4 (green) and FoxP3 (red) expression within CD20 (cyan) and Ki67 (yellow) double positive areas. Original magnification 40x. Scale bars are 100um, except for the zoomed in images corresponding to the white dotted enclosures which are 20um. Box plots showing the normalized per imaged area numbers of FOXP3^hi^ CD4 T cells within B cell follicles, as quantified by Histocytometry (***p=0.00069, ANOVA). F) Box plots of CD25^hi^FoxP3^hi^ and IL-10^hi^CD25^hi^ FoxP3^hi^ CD4 T cells as measured by Histocytometry. Each blue and red circle represents a single follicle, and data presented are for all follicles measured in NNs (blue) and Ns (red) respectively except for extra-follicular and whole tissue data where each participant tissue is represented by a circle of a different color. ****p<0.0001; ****p<0.0001 (Mann-Whitney U test).

### Cross-neutralization is associated with a lower cell density of IL-10^hi^CD25^hi^FoxP3^hi^ follicular regulatory CD4 T cells

Our transcriptomic analysis revealed a higher frequency of T_REGS_ (defined as FoxP3^hi^ cells) in N compared to NN tissues (**Fig. 3B**). We then used an *in-situ* approach to quantify T_REGS_ in extrafollicular and follicular (T_FR_ cells) LN areas. First, we found a significantly higher number of normalized FoxP3^hi^ CD4 T cells within the follicular areas in Ns as compared to NNs (p=0.00069) **(Fig. 3E, Fig. S3A and S4D).** We further investigated the FoxP3^hi^ CD4 T cell compartment using CD25 (*47*) and IL-10, a cytokine which correlates with increased T_REG_ functionality and suppressive capacity (*48*) **(Fig. S5A).** Analysis of CD25^hi^FoxP3^hi^ CD4 T cells revealed no significant differences in the normalized numbers of extrafollicular CD25^hi^FoxP3^hi^ CD4 T cells amongst the two categories **(Fig.3F).** However, there was a significantly lower number of follicular CD25^hi^FoxP3^hi^ CD4 T cells in Ns compared to NNs tissues was observed (p<0.0001) **(Fig.3F)**. A similar profile was found when the prevalence of follicular IL-10^hi^CD25^hi^FoxP3^hi^ CD4 T cell was calculated (p<0.0001) **(Fig. 3G)**. Our data suggest a GC-specific rather than global effect of increased numbers of functional, IL-10 producing T_FR_ cells in the follicles of NNs which may contribute to limited GC TFH cell responses.

### Cross-neutralization is associated with a higher degree of viral evolution

To explore relationships between the aforementioned CD4 T cell profiles and virus dynamics, we compared NN and N groups according to plasma viral load, the prevalence of HIV RNA^+^ cells and HIV expression level in these cells quantified by *in situ* hybridization, and the intra-host diversity of HIV *env* gene sequences. Plasma viremia was not significantly different between NNs and Ns nor did it correlate with breadth of neutralizing antibodies **(Fig. 4A).** When the *in-situ* expression of HIV mRNA was analyzed, we found significantly more HIV RNA+ cells (p=0.0232) and a trend for higher levels of total virus RNA (p= 0.0704) in the NNs compared to Ns LNs (**Fig. 4B**). Furthermore, there was a positive correlation for FDC area and total FDC-bound RNA virions in both NNs (R^2^=0.9725, p<0.0001) and Ns (R^2^=0.8874, p<0.0001) (**Fig. 4C**). We detected a higher intra-host *env* diversity in neutralizers than in non-neutralizers, based on entropy analysis using high-throughput, single-genome amplification and sequencing (HT-SGS) (*49*) **(Fig. 4D)**. Intra-host diversity of *env* sequences was especially high in one individual with a broad antibody neutralizing profile **(Fig. 4D).** Of note, although this association could be consistent with positive selection by *env*-specific antibodies in neutralizers, we also observed higher intra-host *env* sequence entropy in neutralizers when considering only synonymous variation **(Fig. S5B)**. Moreover, several of the *env* gene regions with particularly high entropy in neutralizers did not correspond to known bNAb epitopes **(Fig. 4D).** Taken together, these findings suggest higher cumulative levels of HIV replication/evolution in neutralizers than in non-neutralizers.

**Fig. 4:**
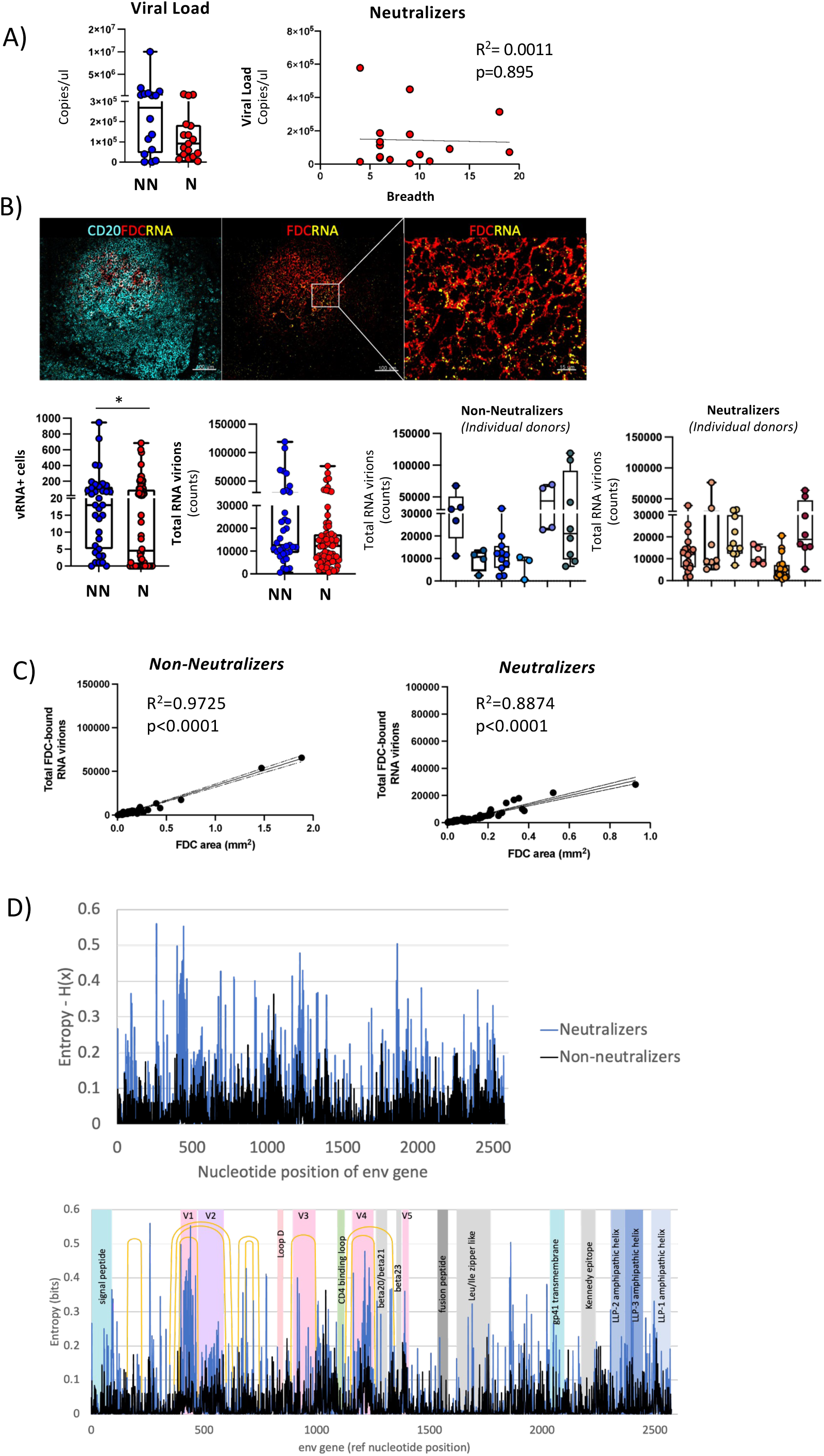
Higher viral evolution is associated with lower *in situ* prevalence of actively transcribed virus in cross-neutralization participants. A) Plasma viral loads (VLs) in all study participants and association of VLs with neutralization breadth (number of isolates recognized out of the 20 isolates tested). B) Representative confocal images showing the distribution of viral RNA (yellow) on FDC networks (red) within B-cell follicles (CD20: cyan) and box plots summarizing the number of RNA+ cells and vRNA+ virions as measured by RNAscope *in situ* hybridization, Histocytometry (cells) and spot analysis (virions) in NNs and Ns. Images were captured at 40x magnification, with 1% zoom and scale is 100um and 15um (zoomed in detail). Each blue and red circle represents a single follicle, and data presented are for all follicles measured in NNs (blue) and Ns (red) tissues *p=0.0232 (Mann-Whitney U test); **p=0.0704 (ANOVA). Graphs showing the number of total RNA virions (counts) in individual NN and N participants are also shown. Each color represents a different participant tissue and individual circles the B-cell follicles in that tissue. C) Association of total RNA virion burden on FDCs and total FDC area (mm^2^) as measured by surface analysis. D) Entropy-H(x) plots showing the location of positions with residue mismatches on alignment to the reference Env HIV sequence, and the distribution of those mismatches over different HIV structural regions. Color shades denote distinct structural regions, and orange arcs the locations of gp120 variable loops.

### Cross-neutralizing activity is associated with a higher in situ prevalence of germinal center B cells expressing a molecular profile favoring the GC development

Finally, we sought to assess whether HIV-1 neutralization could be linked to a specific GC-B cell profile. To this end, we analyzed the numbers of GC-B cells and total B cells (CD20^hi/dim^) *in situ*. Although the total numbers of B cells did not differ between the two groups, LNs of Ns had significantly higher normalized numbers (p = 0.0185) of CD20^hi^Ki67^+^Bcl-6^hi^ B cells than those of NNs **(Fig. 5A** and **Figure S6A)**. Furthermore, significantly higher TFH /B cell (p<0.0001) and a trend for higher Bcl-6^hi^ TFH /Bcl-6^hi^Ki67^hi^ B cells ratios were found in N compared to NNs **(Fig. S6B).** We also performed sc-RNAseq on B cells to elucidate their transcriptomic profile. Analysis of the expression of relevant genes showed a high heterogeneity (23 clusters were identified) of the B cell compartment in our cohort (**Fig.S6C**). Further categorization of B cell subsets revealed significantly higher GC B cell and lower naïve B cell frequencies in Ns than NNs LNs (**Fig. 5B**). This was further supported by the higher expression of CXCR5 and CXCR4 in N than NN LNs **(Fig. 5C)**. Moreover, we found increased expression of factors promoting the GC B cell differentiation (BACH1, STAT3/6), as well as the development of DZ B cell compartment (FOXO1, TGFB/SMAD3/4) and the response to local osmotic stress (NFAT5) in Ns compared to NNs B cells (**Fig. 5C**). NN B cells were found to upregulate type I and type II IFN and mTOR signaling, which could have a detrimental effect on GC B cell development (**Fig. 5C**). Together, these data show that neutralizers are programmed for the development and maintenance of GC B cells. Furthermore, possible molecular targets that could mediate the interaction between GC B cells and TFH cells were investigated across all 8 participants using the NicheNet package in R. Our analysis revealed novel putative ‘molecular-pairs’ that could mediate such interaction, including macrophage migration inhibitory factor-MIF (ligand for CXCR4 and CD74 and potential regulator of B cells at trafficking, survival and antigen presentation level (*50*)) / Integrin beta 8-ITGB8 (unit of the avb8 integrin, a critical activator of TGF-b (*51*)), HMGB2 (critical regulator of V(D)J recombination (*52*)) / LEF1(regulator of TFH cell differentiation (*53*)) and TGF-b1/SMAD3. Therefore, our analysis provides molecules that could serve as targets for in vivo interventions aiming to strengthen the TFH-B cell interaction and bnAb development.

**Fig. 5:**
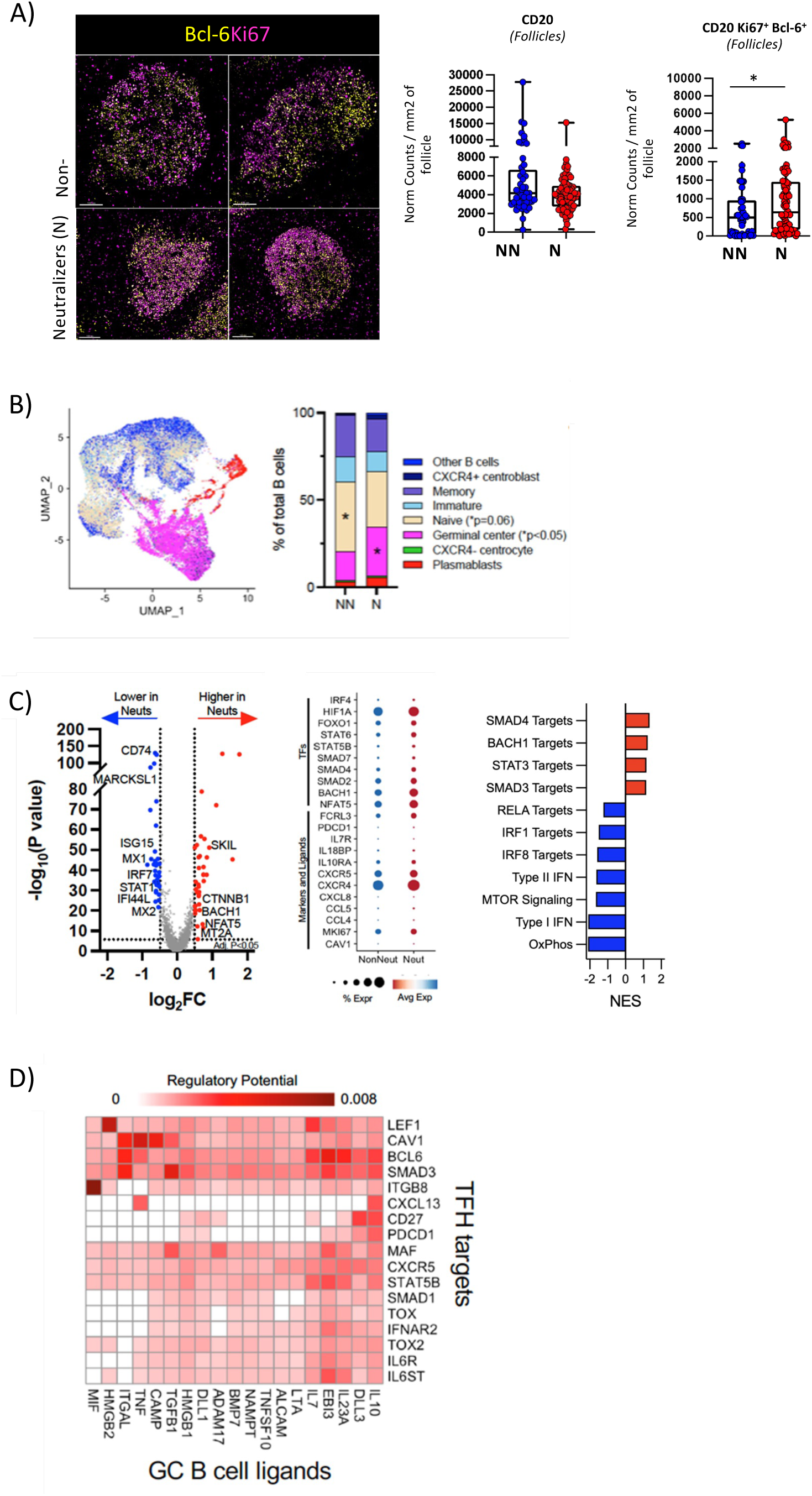
Germinal center B cells express a molecular profile favoring the GC development in cross-neutralization LNs. A) Representative confocal images showing the distribution of Bcl-6 (yellow) and Ki67 (magenta) within the GCs of NNs and Ns. Images were captured at 40x magnification, with 1% zoom and scale is 100um. Box plots and graphs show the pooled normalized numbers of CD20^hi^ and Bcl-6^hi^Ki67^hi^CD20^hi^ B cells as measured by Histocytometry. Circles represent single follicles, and data are from all follicles measured in NNs (blue) and Ns (red) tissues (p=0.1692 and *p=0.0185, ANOVA). B) UMAP projections of B-cell populations in total LMNCs, color-coded by B cell category. C) Volcano plots and dot plots of differentially expressed genes (DEGs) and normalized enrichment scores (NES) of GSEA for select B-cell specific ligands and transcription factors are shown. The DEGs were calculated using the MAST (Hurdle model based) test via the FindMarkers function using the Seurat package in R. The p-values were corrected using the Benjamini-Hochberg method and genes meeting an adjusted p< 0.05 and abs(log_2_FC)>0.5 are highlighted on the volcano plot. Dots in volcano plots represent genes that are significantly downregulated (blue) or upregulated (red) in neutralizers. D) Heatmap of pairwise TFH-GC B cell interactions as identified through ligand-target analysis. Darker colors denote interactions corresponding to an increased regulatory potential.

## Discussion

Development of neutralizing antibodies through vaccination is critical for effective control of viruses. A subset of PLWH, independent of vaccination, can develop neutralizing antibodies against HIV (*54*). Although the characteristics of a variety of bNAbs have been previous explored, the human follicular/GC immune dynamics that mediate the development of these antibodies are largely unknown. In this study, we used LN samples from a well-characterized cohort of ART-naïve people living with chronic HIV to investigate the tissue determinants and immunological characteristics associated with the development of HIV neutralizing antibodies.

HIV is associated with a wide spectrum of lymphadenopathies, including follicular hyperplasia, lysis and involution (*4, 55, 56*). Our morphological analyses showed that Ns LNs are characterised by better preserved follicular/GC areas (smaller follicular areas associated with higher of GC/F and FDC/CD20 area ratios compared to NNs LNs), providing the ground for more efficient development of GC immunoreactions. A recent study has identified a critical role for FoxP3^hi^ TFH cells in regulating follicular size and lifespan(*57*). Interestingly, significantly higher numbers of follicular FoxP3^hi^ CD4 T cells were found in the N compared to NN participants, suggesting that such mechanism may be responsible, at least in part, for the better-regulated follicular development of PLWH with superior neutralizing antibodies.

Differentiation of TFH cells is a multiphase process driven by changes in gene expression due to intrinsic and surrounding microenvironment cues that yield a cell pool with high heterogeneity(*35*). Our flow cytometry analysis revealed a skewed differentiation of bulk CD4 T cells towards the CD27^hi^CD45RO^hi^ memory compartment which is associated with a higher frequency of TFH cells in the N compared to NN participants. CD57^hi^PD1^hi^ human TFH cells have an augmented capacity to secrete IL-4, at least, *in vitro* (*35*), presumably representing Th2-like TFH cells. While there was a significant expansion of Th1-like (CD57^lo^CXCR3^hi^) TFH cells in NN LNs, a cluster signature encompassing Th2-like (CD57^hi^CXCR3^lo^) TFH cells appeared more elevated in Ns (*58*). Furthermore, a significant accumulation of Bcl-6^hi^ TFH cells was observed in the N LNs by *in situ* analysis, which was associated with a significantly higher breadth of neutralizing antibodies. Therefore, neutralizing antibody development in PLWH could be associated with a significant expansion of effector TFH cells that can adopt a Th2-like phenotype, in line with our previously published data regarding the profile of SHIV Env-specific TFH cells in neutralizing non-human primates (*59*).

Consistent with the flow cytometry data, scRNAseq analysis showed an increase in TFH subsets along with a significant decrease in the naïve CD4 T cell compartment in N compared to NN participants. TFH cells from Ns LNs were characterized by significant upregulation of genes favoring their i) development (e.g. IL6ST, GITR, STAT3, Tox2 and Maf (*60, 61*)), ii) function (CXCL13) and iii) interaction with surrounding cells (PDCD1, TIGIT, ICOS). Furthermore, the pathway/targets analysis revealed a significant upregulation of SMAD3/4 and STAT3 targets, indicating a higher operation of TGF-β-induced signaling (*62*), a positive regulator of TFH cell differentiation (*63*), in TFH cells from N participants. This is further supported by the significant increase of the ITGB8 gene transcript in TFH cells in Ns, which encodes the b8 subunit of the αvβ8 integrin, a major positive regulator of TGF-β bioavailability (*64*). Wnt signaling and Tcf7 targets, pathways associated with T cell stemness (*65*), were also significantly upregulated in N compared to NN participants. This profile was associated with significantly higher expression of CD27, a survival factor for memory CD4 T cells and necessary for long-lived T cells (*66*). Altogether, our data indicate that neutralization activity in PLWH is associated with a molecular program favoring the development of TFH cells with increased capacity for self-renewal and longevity.

T_FR_ cells are specialized CD4 T cells that localize in GCs and regulate TFH function by limiting their numbers and curtailing their ability to provide B help through ICOS co-stimulation as well as IL-21 and IL-4 production (*24, 47, 67*).Since T_FR_ cells have been shown to expand both proportionally and numerically during HIV and SIV infection (*24*), we asked if differences in neutralization could also be due to differences in the frequencies of T_REG_ and T_FR_ cell populations between the two study groups. Although significantly higher numbers of follicular FoxP3^hi^ CD4 T cells were found in N participants, the opposite was observed when CD25^hi^FoxP3^hi^ and IL-10^hi^CD25^hi^FoxP3^hi^ follicular CD4 T cells were analyzed. This suggests that TFH cells in N participants are exposed to a less immunosuppressive microenvironment. On the other hand, T_REGS_ exposed to local inflammatory microenvironment adopt a strong suppressive function (*68*). The significant reduction in IL-10^hi^CD25^hi^FoxP3^hi^ T_FR_ cells found in N compared to NN participants further corroborate a less inflammatory follicular environment in these individuals. Further investigation of the mechanism(s) associated with the reduced presence of T_FR_ cells in Ns could yield critical information for the development of broadly neutralizing antibodies.

We observed higher evolution of Env protein in N than NN participants, which corroborates published findings(*69*). This profile, however, was associated with different *in situ* virion dynamics between the two groups. A higher *in situ* viral active transcription was found in NN tissues as evidenced by the number of HIV mRNA+ cells and a trend for higher overall numbers of total virions. The FDC network is a stable ‘reservoir’ of highly infectious virus (*70, 71*). We found a positive association between FDC area and FDC-bound virions both in NN and N tissues, indicating that differences in FDC-mediated viral dissemination(*71, 72*) alone cannot explain the higher levels of viral replication seen in NNs.

With respect to the B cell compartment, our scRNAseq analysis showed a significantly higher prevalence of GC B cells in N participants. B cells from N participants were characterized by a significant upregulation of genes which encode molecules involved in the regulation of B cell differentiation and antibody responses (e.g. BACH1, STAT3)(*73–75*), B cell trafficking (CXCR5, CXCR4)(*76*) as well as protection of GC B cells (NFAT5)(*77, 78*) from the high osmotic stress present in follicular areas. Furthermore, the transcriptional profiling indicates a higher prevalence of DZ GC B cells and preservation of an anatomically distinct DZ (upregulation of FOXO1, CXCR4) in N compared to NN participants where LZ GC B cells may be more dominant (higher expression of mTOR) (*76, 79–81*). Our data revealed increased type I and II IFN signaling signatures in both TFH and B cells in NN participants. Type I IFN signaling could alter the differentiation of CD4 T cells towards a Th1 phenotype (*82*), in line with the significant reduction in numbers of Bcl-6^hi^ TFH cells found in NN participants. On the other hand, it is well established that type I and type II IFNs are potent activators of the mTOR/PI3K pathway (*83*). During acute viral infections, an early boost of both type I and II IFN-mediated signalling followed by PI3K/Akt/mTOR pathway activation is beneficial for GC-formation and for the generation of antigen-specific plasma cells (*84–86*). However, hyperactivation of mTORC1 complex, fuelled by sustained IFN signalling or other stimuli during chronic viral infection, restricts GC-B cells in the DZ which results in decreased access to antigen and TFH cells in the LZ (*85*). Moreover, activation of mTORC1 can induce class switch recombination in murine B cells, favouring IgG1-expressing B cell clones (*86, 87*). Considering that the IgG1 isotype of some bNAbs was reported to exhibit reduced neutralization capacity of viral escape HIV variants compared with IgG3 and IgA1 (*88*), we propose that sustained mTORc1 signalling might be a limiting factor for the generation of potent neutralizing Abs.

Together, our study suggests that the development of neutralizing antibodies during chronic HIV infection is multifactorial, and associated with i) a CD4 T cell programing favoring the differentiation towards TFH cells, especially ones with increased stemness and long-lived capacity and decreased prevalence of a Th1-like TFH phenotype, ii) a B cell programing enabling the formation of anatomical DZ and DZ GC B cells, iii) reduced GC suppressive activity and iv) higher evolution of Env. One limitation in our clinical study, is that the exact timing of participant HIV-1 acquisition could not be ascertained. As such, differences in the approximated lengths of viral infection could exist. Even with this limitation however, the requirement for a better-preserved follicular microarchitecture remains as we found i) several predicted classes of bNAbs in the plasma of participants categorized as “neutralizers” and ii) a strong correlation between the observed TFH phenotypes and the development of breadth. We thus propose that a preserved follicular microarchitecture provides the ground for the orchestrated action of the aforementioned cell phenotypes resulting in the development of neutralizing antibodies. Our analysis also elucidates the B and T cell molecules that could represent novel targets for innervations aiming to strength the TFH-B cell interaction and antibody responses.

## Materials and Methods

### Study Cohort and Ethics Statement

Plasma samples, lymph node cell suspensions and biopsies were obtained from untreated, chronically infected PLWH. All tissue samples from PLWH were procured with explicit written informed consent from participants prior to donation, adhering strictly to the principles outlined in the Declaration of Helsinki. The utilization of these samples was formally sanctioned by both the Research Committee and the Ethics in Research Committee of the National Institute of Respiratory Diseases “Ismael Cosío Villegas,” Mexico City as part of the “C71-18” protocol. None of the participants included in the protocol had active opportunistic infections and all were HBV and HCV negative. Lymph node (LN) biopsies were obtained from PLWH who had palpable LN in the cervical area or LN detected by ultrasound in the inguinal area. The control tissue samples were retrieved from the archives of the Institute of Pathology of Lausanne University Hospital, Switzerland and their use was approved by the Ethical Committee of the Canton de Vaud, Switzerland (protocol number 2021-01161). Anonymized, discarded pathologic tonsil specimens were obtained from Children’s National Medical Center (CNMC) under the auspices of the Basic Science Core of the District of Columbia Developmental Center for AIDS Research. The CNMC Institutional Review Board determined that study of anonymized discarded tissues did not constitute ‘human subjects research’. Sample sizes were not predetermined by power calculations, and investigators were not blinded to group identity during the study. Plasma viral load (pVL) was quantified by automated real-time PCR using the m2000 system (Abbott, Abbott Park, IL, USA). The range of detection for pVL was 40 to 10,000,000 copies/ml. CD4+ T cell counts were determined by flow cytometry using the TruCount kit in a FACSCanto II instrument (BD Bioscience, San Jose, CA, USA), according to the manufacturer’s instructions. Tonsilar and LN cell suspensions were stored at liquid nitrogen until further use.

### Flow Cytometry-data acquisition and analysis

Surface and intracellular staining of LN and tonsil derived cells was carried out using appropriate, titrated antibodies (**Table S3**). Events were collected on a BD X50 Symphony (BD Biosciences), and electronic compensation was performed with antibody capture beads (BD Biosciences). Data were analyzed using FlowJo version 10.8.1 (TreeStar Inc, BD). Dimensionality reduction analysis was performed using the t-SNE and FlowSOM plugins on FlowJo (TFH populations). For t-stochastic neighborhood embedding (t-SNE) analysis of TFH, individual populations were manually gated in FlowJo, and fcs files were exported for each participant. Files were pre-processed to exclude batch-to-batch variations, and concatenated to a single file down-sampled to 11481 cells/sample, yielding a file suitable for t-SNE analysis. t-SNE parameters were set to 1000 iterations, learning rate: 2818 and perplexity 30. Clustering analysis using the FlowSOM plugin (FlowJo) was performed for the following 11 markers: TIGIT, ICOS, CD95, CD57, CXCR3, OX40, CTLA-4, PDL-1, CD226, OX40L, TIM-3, for a grid size of 10x 10, yielding 8 meta-clusters. For visualization, clustering results were overlayed on the t-SNE projection using the ClusterExplorer plugin. Differences in frequencies in FlowSOM generated clusters were calculated using an unpaired t-test in GraphPad Prism version 9.3.1.

### Multiparameter Confocal Imaging-data acquisition and analysis

Unconjugated primary and secondary, and conjugated antibodies were used for the staining of paraformaldehyde-fixed, paraffin embedded tissue sections **(Table S4).** For virus RNA visualization, RNAscope in situ hybridization was performed according to the manufacturer’s instructions for formalin-fixed paraffin embedded tissues using dedicated HIV RNA probes (Cat No. 416111, Advanced Cell Diagnostics, Hayward CA) and the RNAscope Multiplex Fluorescent Reagent Kit v2 (Advanced Cell Diagnostics, Hayward CA). The assay was performed as previously reported (*89*) but without proteinase K treatment in order to preserve the human CD20 epitope. Confocal images were obtained on a Leica TCS SP8 confocal system, at 512 x 512-pixel density and 1x optical zoom using a 40x objective (NA 1.3), unless otherwise stated. No frame averaging or summing was used while obtaining the images. To ensure accurate representation and minimize selection bias, at least 50% of the tissue was imaged. Fluorophore spillover, when present, was corrected by imaging tissues stained with single antibody-fluorophore combinations, and by creating a compensation matrix via the Leica LAS-AF Channel Dye Separation module (Leica Microsystems) per user’s manual. Confocal images were analyzed with the ImageJ software (*90*) and Imaris version 9.5.0 (Bitplane). Histocytometry analysis was performed to generate quantitative data from the images, as previously described (*29, 35, 89*).Fragmented follicles, follicles partial on edge and follicles with indeterminate boundaries were excluded from the analysis. The number of FDC-bound virions in RNAscope analysis was also measured with the software Imaris. Quantitation was performed using the Spots creation feature after appropriately masking the vRNA+ channel to reveal colocalization with FDC surfaces. For RNA virion quantification, a thresholding diameter of 2um was used consistent with previous work in our lab (*91*).

### Single cell RNA sequencing (scRNAseq)-data analysis

FASTQ files were uploaded to Cell Ranger on the 10X Genomics cloud, and no depth normalization was carried out. The filtered count matrix was then analyzed using the Seurat package in R. Cell annotation was carried out using SingleR and the reference expression dataset was derived from the MonacoImmuneData atlas from the celldex R package. Doublet cells were removed from the analysis using the DoubletFinder package in R. Differentiation gene expression was assessed using the MAST R package. Clustered cells were visualized using Uniform Manifold Approximation and Projection (UMAP), and global differences between clusters were assessed using Principal component analysis (PCA).

### High-throughput, single-genome amplification and sequencing (HT-SGS) of HIV *env*

Plasma samples were thawed at room temperature, followed by a brief centrifugation to collect liquid from the sides of tubes. RNA extraction from virions in the supernatant was conducted using the QIAamp Viral RNA Mini Kit (Qiagen, 52906), following the instructions provided by the manufacturer. Subsequence processing steps were similar to those described in our previous study(*49*).

### Statistics

We compared differences between groups (non-neutralizers, neutralizers) in flow cytometry, Histocytometry, and sc-RNA using an ANOVA. A parametric t-test was used for the t-SNE cluster analysis whereas data with more than one explanatory variable were analyzed using a two-way ANOVA followed by post-hoc comparisons (Tukey). In those cases where the dependent variable was not normally distributed according to the Lilliefors and/or Shapiro-Wilk normality tests, a box-cox transformation was applied, and when the transformation did not render a normally distributed variable, the non-parametric Mann-Whitney U test for unpaired variables was used. GraphPad Prism Version (10.2.1), JASP (version 0.18.3) and the program R studio (Version 2023.06.2+561 with packages: nortest (10.1-4) and ggplot2 (3.5.0) were used to create the graphs and perform the t-tests, Mann-Whitney U tests and ANOVA tests respectively. All quantitative data show means (±*SD*) except from box plots which show inter-quartile ranges (IQR). P values and probability values of less than 0.05 were considered statistically significant. Statistical significance p-value key is the following: * = p value <0.05, ** = p value < 0.01, *** = p value <0.001, **** = p value <0.0001.

## Supplementary Materials

Materials and Methods

**Fig.S1**: Cross-neutralization profiles of NN and N study participants, and heterogeneity of B-cell follicle shape.

**Fig.S2**: Phenotypic characterization of TFH cells.

**Fig.S3**: *In situ* characterization of TFH cells.

**Fig.S4**: Identification of CD4 T cell subsets using sc-RNA analysis of LMNCs.

**Fig.S5:** Regulatory T cell HistoCytometry gating strategy and Entropy-H(x) plots

**Fig.S6**: Identification of B cell clusters using sc-RNA analysis of LMNCs.

**Table S1:** Clinical Information of study participants

**Table S2:** Study Assays

**Table S3:** Flow Cytometry Antibodies

**Table S4:** Multiparameter Imaging Antibodies

## Acknowledgements

The authors would like to thank Dr Natalie Piazzon (operational director of the Tissue Biobank) and Damien Maison and Emilie Lingre, Institute of Pathology, CHUV, for their help with the tissue processing.

## Author Contributions

E.M. and A.A.S. performed experiments, analyzed and interpreted data and drafted the manuscript; S.O.D., S.G., P.M.D.R.E., M.H.B., S.H.K., F.B., M.O. and C.M.B. performed experiments, A.B.E. participated in sc-RNA analysis and drafted the manuscript, N.D.R. and E.B. analyzed and interpreted data and drafted the manuscript, F.T.R., S.A.R., G.R.T. and L.de L. provided participant material and clinical data, S.M., J.R.M., R.P.S, R.A.K. provided supervision, interpreted the data and critically revised the manuscript C.P conceived, designed and supervised the study, interpreted data, and revised the manuscript. All authors have read and approved the final version for submission.

## Data sharing statement

The authors agree to share all publication-related data. For further information, please contact the corresponding author at

