## Supplementary information for "Neutralization activity in chronic HIV infection is characterized by a distinct programming of follicular helper CD4 T cells"

### Supplemental Materials and Methods

#### Sample Collection and Processing

Upon receipt of tissue, lymph nodes and tonsils were washed with ice-cold medium R-10 (RPMI 1640 supplemented with 10% fetal bovine serum, 2 mM L-glutamine, 100 U/mL penicillin and 100 µg/mL streptomycin (Invitrogen). All LN were tested for the presence of fungi and bacteria by culture and for the presence of *Mycobacterium tuberculosis* by the GenXpert assay (Cepheid Inc). Histopathological studies were performed to discard any neoplasia. After receipt, LN biopsy tissue samples were cleaned of surrounding fatty tissue and cut in three small pieces. The first one was used for microbiological studies, the second one was placed in freshly prepared paraformaldehyde (4% in PBS) for 24 hours and then transferred to Ethanol (EtOH) 80% to follow the procedure for paraffin inclusion. The third piece was placed in R10 media and immediately processed to obtain lymph node mononuclear cells (LNMC) by manual disruption using a 70um mesh and a syringe embolus. Obtained cells were cryogenically preserved until utilized for subsequent experiments. A similar protocol was used for tonsil FFPE preparation, only tissues were placed in 10% Neutral Buffered Formalin for 24hrs before being embedded in paraffin and sectioned for multiplex confocal microscopy analysis. Tonsil LNMC were prepared in the same way as LN LNMCs.

#### Flow cytometry

Lymph node mononuclear cells (1-2x$10^6$ cells/mL) from HIV+ lymph nodes and HIV- tonsils were thawed, rested for 1-hour at 37$^o$C, incubated with the Blue or Aqua LIVE/DEAD Fixable Dead Cell Stain Dye (Invitrogen), and surface-stained with titrated amounts of fluorophore-conjugated antibodies against: CD4, CD27, CD45RO, PD-1, CXCR5, TIGIT, CD95, CXCR3, ICOS, PDL-1, CD226, CD57, OX40, CTLA-4, OX40L, TIM3, CD8, CD19, CD130 for T cell phenotyping

(**Table S3**). For intracellular staining, cells were washed with PBS, fixed, permeabilized using the BD Transcription Factor Buffer Set (BD Biosciences), and stained with titrated amounts of fluorophore-conjugated antibodies against: AHR, CAV-1 and TCF1. Cells were then washed and fixed with 2% paraformaldehyde.

**Multiparameter Confocal Imaging**

Paraffin tissue sections 7-10 micron thick were deparaffinized and rehydrated through sequential Xylene-Ethanol-di$H_2$0 solutions. Heat Induced Epitope Retrieval was performed using the Borg Decloaker reagent (Biocare Medical) for 15 minutes at 110$^o$C in a chamber pressurized at 5-6 PSI. This was followed by a blocking and permeabilization step using a Phosphate Buffered (PBS)/ Bovine Serum Albumin (BSA)/ Triton-X solution, after which tissue sections were stained overnight at 4$^o$C with titrated amounts of primary antibodies. The next day, tissue sections were washed with PBS, and stained with secondary fluorophore-conjugated antibodies, followed by conjugated antibodies **(Table S4)**. At the end of all staining rounds, a final washing step with PBS was performed, and the nuclear stain JOJO-1 or JOPRO-1 was applied (ThermoFisher Scientific). Slides were mounted with a glass coverslip using Fluoromount G (Southern Biotech). For RNAscope assays, sections were baked for 1 hour at 60$^o$C and deparaffinized using serial xylene and ethanol baths. This was followed by an antigen retrieval step at 110$^o$C for 15 minutes. Subsequently, sections were incubated with the HIV-1 Clade B-specific probes at 40$^o$C for 2 hours in a HybEz hybridization oven (ACD) before being subjected to 4 rounds of signal amplification performed using the RNAscope amplification reagents. To visualize the RNA signal, a tyramide-based detection system was used (TSA Plus Cyanine 5, Akoya Biosciences) per manufacturer's instructions. All additional immunofluorescent stainings, as applicable, were performed at the end of the RNAscope protocol.

### Imaging data analysis

*ROI morphological analysis*: to avoid methodological bias two different approaches were applied for the measurement of total surface areas in regions of interest (ROI: Follicular Areas). First, ROIs were analyzed with the Image J software. All images were transformed to binary and two univariate morphological features were measured for each ROI: Surface area (Sa) and Circularity. Then, Surface area was transformed/normalized based on the scale bar of each binary image by using the formula $Sa_{NORM}$ = Sa*Pixels of scale bar/Length of scale bar, in order to be able to compare ROIs obtained from different tissues. Next, trends were compared with those obtained from Imaris with the Surface Creation module, to ensure the two methods yielded comparable results before undertaking further within-ROI quantifications with Imaris.

*Histocytometry analysis*: briefly, 3-dimensional segmented surfaces (based on nuclear signal) of background and spillover corrected images were generated with Imaris via the Surface Creation module. Average voxel intensities for all channels, as well as volume and sphericity of the 3-dimensional surfaces generated from histocytometry were exported in Microsoft Excel format. After conversion of the files to comma separated value (.CVS) files, the data were imported into FlowJo (version 10) for further analysis/quantification. Analysis included well-defined areas devoid of background staining and data were quantified either as relative frequencies or as cell counts normalized to total follicular area ($mm^2$) screened.

### Neutralization Assays

The neutralization activity of plasma was measured by incubating 10µl of five-fold serially diluted serum in cDMEM with 40ul of diluted HIV-1 Env-pseudotyped virus for 30 minutes at 37$^{o}$C in a 96-well CulturPlate (Perkin Elmer). 20 µl of TZM-bl cells (10,000 cells/well) with or without 70µg/ml DEAE-Dextran was then added and incubated overnight at 37$^{o}$C. Each experiment plate also had a column of cells only (no Ab or virus) and a column of virus only (no Ab) as controls for background TZM-bl luciferase activity and maximal viral entry, respectively. Serial dilutions were performed with a change of tips at each dilution step to prevent carryover. The following day, all wells received 100µl of fresh cDMEM and were incubated overnight at 37°C. The following day, 50µl of Steadylite Plus Reporter Gene Assay System (PerkinElmer) was added to all wells, and plates were shaken at 600RPM for 15 minutes. Luminometry was then performed on a SpectraMax L (Molecular Devices) luminometer. Percent neutralization is determined by calculating the difference in average RLU between virus only wells (cells + virus column) and test wells (cells + plasma/Ab sample + virus), dividing this result by the average RLU of virus only wells (cell + virus column) and multiplying by 100. Background is subtracted from all test wells using the average RLU from the uninfected control wells (cells only column) before calculating the percent neutralization. Neutralizing plasma or serum antibody titers are expressed as the dilution required to achieve 50% neutralization (ID50) and calculated using a dose-response curve fit with a 5-parameter nonlinear function. For antibody class prediction, the neutralizing activity data were imported into the NFPws Neutralization Fingerprinting Web Server (http://iglab.accre.vanderbilt.edu/NFPws/) and analyzed as previously described [25].

**Single cell RNA sequencing (scRNAseq)**

scRNAseq was performed on LN mononuclear cells from N (n = 4) and NNs (n = 4). The 3' V3 chemistry kit (10X Genomics) was used according to manufacturer's recommendations. Briefly, single cell Gel Beads-in-Emulsion (GEMs) were generated by combining 10,000 LN cells, barcoded Single Cell 3' v3.1 beads, and partitioning oil. This reaction was then run on a 10X Chromium Controller machine (10X Genomics). Afterwards, GEMs were processed to remove gel beads, and cells were lysed and underwent a reverse transcription reaction in a C1000 Touch

Thermal Cycler (BioRad) to generate barcoded cDNA. The GEM mixture was then cleaned using Dynabeads to remove oil and cDNA was amplified. Amplified cDNA was then cleaned using the SPRIselect reagent and the quality/quantity of cDNA was assessed using a 2100 Bioanalyzer Instrument (Agilent Technologies). This cDNA then underwent fragmentation, end repair, A-tailing, and adaptor ligation according to manufacturer protocol recommendations. Library quality was assessed using Bioanalyzer. Libraries were then sent for sequencing at the Beijing Genomics Institute (BGI). Libraries were sequenced on a DNBseq-T7 (MGI Tech) machine targeting 20,000 reads per cell.

**High-throughput, single-genome amplification and sequencing (HT-SGS) of HIV *env***

For reverse-transcription of the virus RNA, SuperScript III (ThermoFisher Scientific, 18080093) was used along with a gene specific reverse transcription primer, which is composed of an RNA binding site, 8 random-base Unique Molecular Identifiers (UMIs) and a reverse primer binding site for PCR (CCCGCGTGGCCTCCTGAATTATNNNNNNNNGTCATTGGTCTTAAA GGTACCTG). After RNA denaturation at 65 °C for 10 min, the RNA was reverse transcribed under following conditions: 50 °C for 10 min, 85 °C for 10 min, and 4°C hold. The cDNA was purified with a 2.2:1 volumetric ratio of RNAClean XP solid phase reverse immobilization beads (A63987, Beckman Coulter). The amount of cDNA was determined through limiting-dilution PCR using fluorescence-assisted clonal amplification (FCA)[89] to quantify cDNA copies. Next, the full-length *env* gene region was amplified in PCR for library preparation using the Advantage 2 PCR kit (Takara Bio, 639206) and forward (GAGCAGAAGACAGTGGCAATGA) and reverse (CCCGCGTGGCCTCCTGAATTAT) primers. The thermocycling conditions were as follows: initial denaturation at 95˚C for 1 min, 31 cycles of denaturation at 95˚C for 10 sec, annealing at 64˚C for 30 sec, and extension at 68˚C for 3 min, and final extension at 68˚C for 10 min. The final concentration of each reagent in PCR was 400 nM of forward and reverse primers, 200 mM of dNTP, 1X of Advantage 2 Buffer and 1X of Advantage 2 Polymerage Mix. Amplified products of approximately 3 kilobases were incorporated into sequencing libraries using the SMRTbell Express Template Prep Kit 2.0 (100-938-900, Pacific Biosciences) and Barcoded Overhang Adapter kit 8A and 8B (101-628-400 and 101-628-500, Pacific Biosciences) for multiplexing targeted sequencing. The libraries were treated by primer annealing and polymerase binding using the Sequel II Binding Kit 2.0 and Int Ctrl 1.0 (101-842-900, Pacific Biosciences), and then sequenced on a Sequel II system (Pacific Biosciences) with a 30-hour movie time under circular consensus sequencing (CCS) mode. Bioinformatic analysis of sequence data was performed using a custom bioinformatic pipeline presented at https://github.com/niaid/UMI-pacbio-pipeline, with modifications to accommodate HIV *env* sequence length and primer sequences. Processed *env* sequences produced by this pipeline were translated and aligned with MAFFT [90] and manually curated. Pal2Nal was used to codon align the cDNA sequences to the protein alignment [91]. Shannon entropy was calculated per position for each position, and also distinguishing synonymous and nonsynonymous positions (positions which could have both synonymous and nonsynonymous changes were not included in the synonymous/non-synonymous groups).

A)

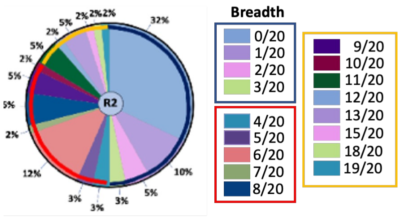

B)

| | A | A | AD | AE | AE | AE | AE | AE | AG | B | B | BC | BC | C | C | C | C | C | D | D |
|---|---|---|---|---|---|---|---|---|---|---|---|---|---|---|---|---|---|---|---|---|
| PID | KER2008.12 | Q259.17 | Q461.e2 | 620345.c1 | CNE5 | CNE55 | M02138 | TH976.17 | 242-14 | 7165.18 | YU2.DG | CH038.12 | CH070.1 | 0013095-2.11 | 001428-2.42 | 26191-2.48 | 3168.V4.C10 | ZM135.10a | 231965.c1 | 247-23 |
| Non neutralizers | (n=16) | | | | | | | | | | | | | | | | | | | |
| HIV #1 | <20 | <20 | <20 | <20 | <20 | <20 | <20 | <20 | <20 | <20 | <20 | 22 | <20 | <20 | <20 | <20 | <20 | <20 | <20 | <20 |
| HIV #2 | <20 | <20 | <20 | <20 | <20 | <20 | <20 | <20 | <20 | <20 | <20 | <20 | <20 | <20 | <20 | <20 | <20 | <20 | 54 | <20 |
| HIV #3 | <20 | <20 | <20 | <20 | <20 | <20 | <20 | 63 | <20 | 28 | <20 | <20 | <20 | <20 | <20 | <20 | <20 | <20 | <20 | <20 |
| HIV #4 | <20 | <20 | <20 | <20 | <20 | <20 | <20 | <20 | <20 | <20 | <20 | <20 | <20 | <20 | <20 | <20 | <20 | <20 | <20 | <20 |
| HIV #5 | <20 | <20 | <20 | <20 | <20 | <20 | <20 | <20 | <20 | <20 | <20 | <20 | <20 | <20 | <20 | <20 | <20 | <20 | <20 | <20 |
| HIV #6 | <20 | <20 | <20 | <20 | <20 | <20 | <20 | <20 | 22 | 64 | <20 | <20 | <20 | <20 | <20 | <20 | <20 | <20 | <20 | <20 |
| HIV #7 | <20 | <20 | <20 | <20 | <20 | <20 | <20 | <20 | 35 | 54 | <20 | <20 | <20 | <20 | <20 | <20 | <20 | 26 | <20 | <20 |
| HIV #8 | <20 | <20 | <20 | <20 | <20 | <20 | <20 | <20 | <20 | <20 | <20 | <20 | <20 | <20 | <20 | <20 | <20 | 26 | <20 | <20 |
| HIV #9 | <20 | <20 | <20 | <20 | <20 | <20 | <20 | <20 | <20 | <20 | <20 | <20 | <20 | <20 | <20 | <20 | <20 | 67 | <20 | <20 |
| HIV #10 | 40 | <20 | <20 | <20 | <20 | <20 | <20 | <20 | <20 | 27 | <20 | <20 | <20 | <20 | <20 | <20 | <20 | 117 | <20 | <20 |
| HIV #11 | <20 | <20 | <20 | <20 | <20 | <20 | <20 | <20 | 26 | 22 | <20 | <20 | 26 | <20 | <20 | <20 | <20 | 23 | <20 | <20 |
| HIV #12 | <20 | <20 | <20 | <20 | <20 | <20 | <20 | <20 | <20 | <20 | <20 | <20 | <20 | <20 | <20 | <20 | <20 | <20 | <20 | <20 |
| HIV #13 | <20 | <20 | <20 | <20 | <20 | <20 | <20 | <20 | 21 | <20 | <20 | <20 | <20 | <20 | <20 | <20 | <20 | <20 | <20 | <20 |
| HIV #14 | <20 | <20 | <20 | <20 | <20 | <20 | <20 | <20 | <20 | <20 | <20 | <20 | <20 | <20 | <20 | <20 | <20 | <20 | <20 | <20 |
| HIV #15 | 27 | <20 | <20 | <20 | <20 | <20 | <20 | <20 | <20 | <20 | <20 | <20 | <20 | <20 | <20 | <20 | <20 | <20 | <20 | <20 |
| HIV #16 | <20 | <20 | <20 | <20 | <20 | <20 | <20 | <20 | <20 | <20 | <20 | <20 | <20 | <20 | <20 | <20 | <20 | 21 | <20 | <20 |
| Neutralizers | (n=17) | | | | | | | | | | | | | | | | | | | |
| HIV #17 | 59 | 42 | <20 | <20 | 32 | <20 | 137 | 146 | 53 | 68 | 40 | 62 | <20 | 185 | 83 | 24 | <20 | 35 | <20 | <20 |
| HIV #18 | <20 | <20 | <20 | <20 | <20 | <20 | 76 | 39 | 75 | <20 | <20 | <20 | <20 | 70 | <20 | <20 | <20 | 53 | <20 | <20 |
| HIV #19 | <20 | <20 | <20 | <20 | <20 | 63 | 464 | 91 | 28 | 45 | 26 | <20 | <20 | 112 | <20 | 21 | <20 | 52 | <20 | <20 |
| HIV #20 | 77 | <20 | <20 | <20 | <20 | <20 | 232 | 180 | 322 | 38 | 73 | <20 | <20 | 90 | 44 | <20 | <20 | 54 | <20 | 82 |
| HIV #21 | <20 | <20 | <20 | <20 | <20 | <20 | 115 | 118 | 51 | 102 | <20 | 32 | <20 | 240 | <20 | <20 | <20 | 82 | <20 | 23 |
| HIV #22 | 32 | 86 | <20 | <20 | <20 | <20 | <20 | 27 | <20 | <20 | 33 | 96 | 150 | 27 | 41 | 60 | <20 | <20 | <20 | 78 |
| HIV #23 | 345 | 76 | 51 | 82 | 58 | 176 | 130 | 54 | 129 | 262 | 129 | 255 | 360 | 179 | 170 | 43 | 54 | 153 | 631 | <20 |
| HIV #24 | 83 | <20 | <20 | 111 | <20 | 175 | 381 | 169 | 137 | 300 | 213 | 102 | <20 | 160 | 35 | 38 | <20 | 328 | 28 | <20 |
| HIV #25 | 76 | 64 | <20 | 52 | <20 | 56 | 105 | 64 | 83 | 70 | 235 | <20 | <20 | 139 | 895 | 28 | 154 | 183 | 20 | <20 |
| HIV #26 | <20 | 161 | <20 | 36 | 27 | 95 | 150 | 97 | 57 | 91 | 135 | <20 | 39 | 177 | 85 | 32 | <20 | 283 | 35 | <20 |
| HIV #27 | 782 | 198 | <20 | 78 | 109 | 107 | 1,046 | 506 | 1,283 | 1,147 | 195 | 94 | 154 | 142 | 537 | 34 | 100 | 230 | 430 | 208 |
| HIV #28 | <20 | 174 | <20 | <20 | <20 | <20 | 42 | 154 | 97 | 114 | <20 | <20 | 76 | <20 | <20 | <20 | 25 | 149 | <20 | <20 |
| HIV #29 | <20 | <20 | 128 | <20 | <20 | <20 | <20 | 54 | 117 | 146 | <20 | <20 | <20 | 288 | <20 | <20 | <20 | 280 | <20 | <20 |
| HIV #30 | <20 | <20 | <20 | <20 | <20 | 31 | <20 | <20 | <20 | 266 | 389 | 496 | 231 | <20 | <20 | <20 | <20 | <20 | <20 | <20 |
| HIV #31 | <20 | <20 | <20 | 381 | 134 | 479 | 545 | 177 | 511 | 37 | 31 | <20 | <20 | 54 | 85 | <20 | <20 | <20 | 662 | 27 |
| HIV #32 | 42 | <20 | <20 | <20 | <20 | 83 | <20 | <20 | 71 | 96 | 145 | <20 | <20 | 177 | 22 | <20 | <20 | <20 | 33 | <20 |
| HIV #33 | 23 | <20 | <20 | <20 | <20 | <20 | <20 | <20 | 72 | 100 | 137 | 47 | <20 | 311 | <20 | <20 | <20 | 74 | <20 | <20 |

C)

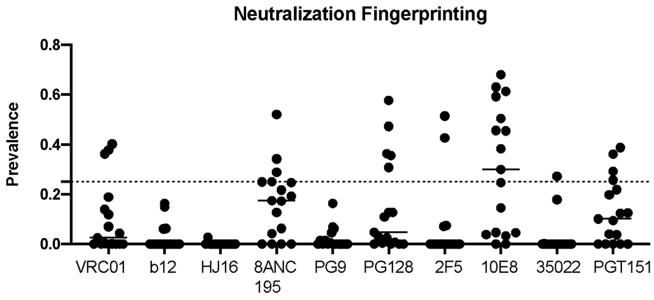

D)

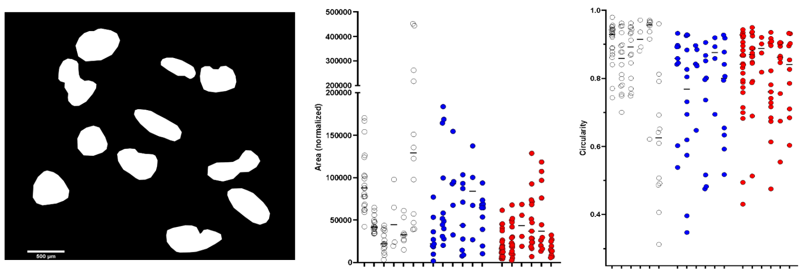

E)

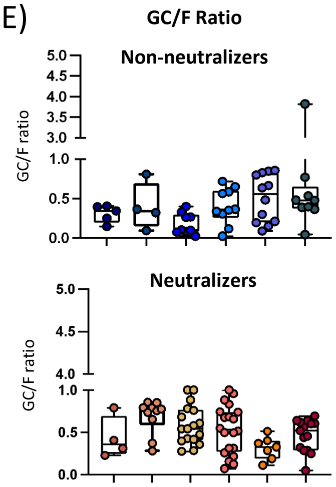

F)

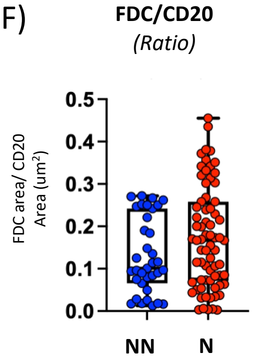

G)

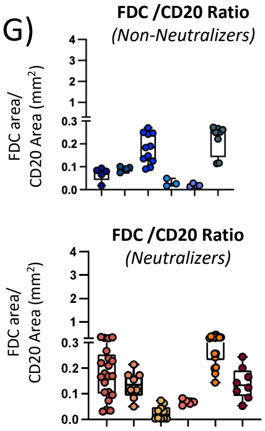

**Fig. S1**: **Cross-neutralization profiles of NN and N study donors, and heterogeneity of B-cell follicle shape**. A) Pie-chart depicting the distribution of cross-neutralization breadth, defined as the percentage of the number of isolates recognized with an $ID_{50}$>40 out of 20 isolates, tested among study participants and B) Profiles of plasma neutralizing activity ($ID_{50}$) in the 33 $HIV^{+}$ donors included in the study color coded by magnitude (green=low, yellow=medium, orange=high). Plasmas were assayed against a panel of 20 different HIV isolates. C) Dot plot showing the distribution of predicted bNAb lineages in the serum of neutralizers as calculated with the NFPws Neutralization Fingerprinting Web Server (http://iglab.accre.vanderbilt.edu/NFPws/). D) Representative example of a binary image used for area determination with ImageJ/Fiji; the areas and circularity in Healthy controls (white circles), Non-neutralizers (blue circles) and Neutralizers (red circles) are shown. Each circle represents an individual follicle in the tissue under study. E-G) Box plots showing the heterogeneity of GC/F ratios in each donor under study, FDC/CD20 ratios in NNs and Ns as calculated using data from pooled quantitative imaging analysis and Histocytometry, as well as the heterogeneity of FDC/CD20 ratios in NNs and Ns as measured by Histocytometry (p=0.144). Circles represent individual follicles. p values represent probabilities as calculated by an ANOVA.

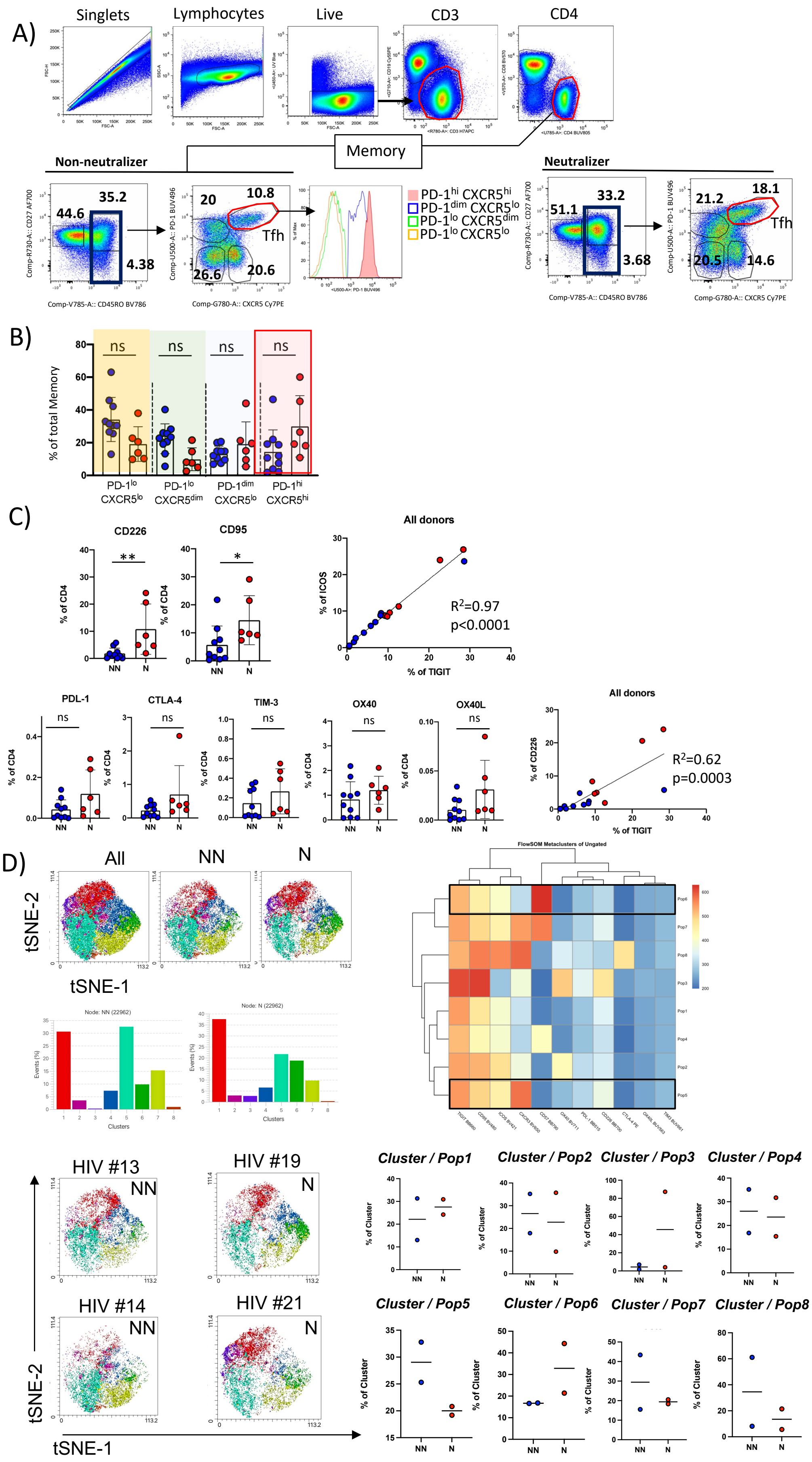
A)
Singlets
Lymphocytes
Live
CD3
CD4
Memory
Non-neutralizer
Neutralizer
35.2
44.6
4.38
20
10.8
Tfh
26.6
20.6
PD-1hi CXCR5hi
PD-1dim CXCR5lo
PD-1lo CXCR5dim
PD-1lo CXCR5lo
33.2
51.1
3.68
21.2
18.1
20.5
14.6
Comp-V785-A:: CD45RO BV786
Comp-G780-A:: CXCR5 Cy7PE
Comp-R730-A:: CD27 AF700
Comp-U500-A:: PD-1 BUV496
B)
% of total Memory
ns
PD-1lo CXCR5lo
PD-1lo CXCR5dim
PD-1dim CXCR5lo
PD-1hi CXCR5hi
C)
CD226
CD95
All donors
% of CD4
% of ICOS
% of TIGIT
% of CD226
R2=0.97
p<0.0001
PDL-1
CTLA-4
TIM-3
OX40
OX40L
R2=0.62
p=0.0003
NN
N
D)
All
NN
N
tSNE-1
tSNE-2
FlowSOM Metaclusters of Ungated
Node: NN (22962)
Node: N (22962)
Events (%)
Clusters
HIV #13
HIV #19
HIV #14
HIV #21
Cluster / Pop1
Cluster / Pop2
Cluster / Pop3
Cluster / Pop4
Cluster / Pop5
Cluster / Pop6
Cluster / Pop7
Cluster / Pop8
% of Cluster

**Fig. S2**: **Phenotypic characterization of $T_{FH}$ cells**. A) Gating strategy for $T_{FH}$ cell characterization in LN cell suspensions using flow cytometry. B) Cumulative flow cytometry frequencies of PD-$1^{lo/dim/hi}$ CXCR5 $^{lo/dim/hi}$ populations within the CD$27^{hi/lo}$ CD45$RO^{hi}$ CD4 T cell memory compartment in NNs and Ns. C) Cumulative flow cytometry frequencies of inhibitory and activating receptors CD226, CD95, PDL-1, CTLA-4, TIM-3, OX40, and OX40L within the PD-$1^{hi}$ CXCR$5^{hi}$ $T_{FH}$ cell compartment in NNs and Ns, and association of TIGIT expression with ICOS and CD226. CD226 **p= 0.0075. CD95 *p=0.0312 (Mann Whitney U test). D) Overlays showing the 8 $T_{FH}$ cell clusters identified through t-SNE (t-stochastic neighbor embedding) analysis of flow cytometry using data from 4 samples with sufficiently representative TFH populations (NN: HIV#13, 14 and N: HIV#19, 21), and their relative representation in the 4 donors analyzed. Clustering was based on the markers TIGIT, ICOS, CD95, CD57, CXCR3, OX40, CTLA-4, PDL-1, CD226, OX40L and TIM-3.

A)

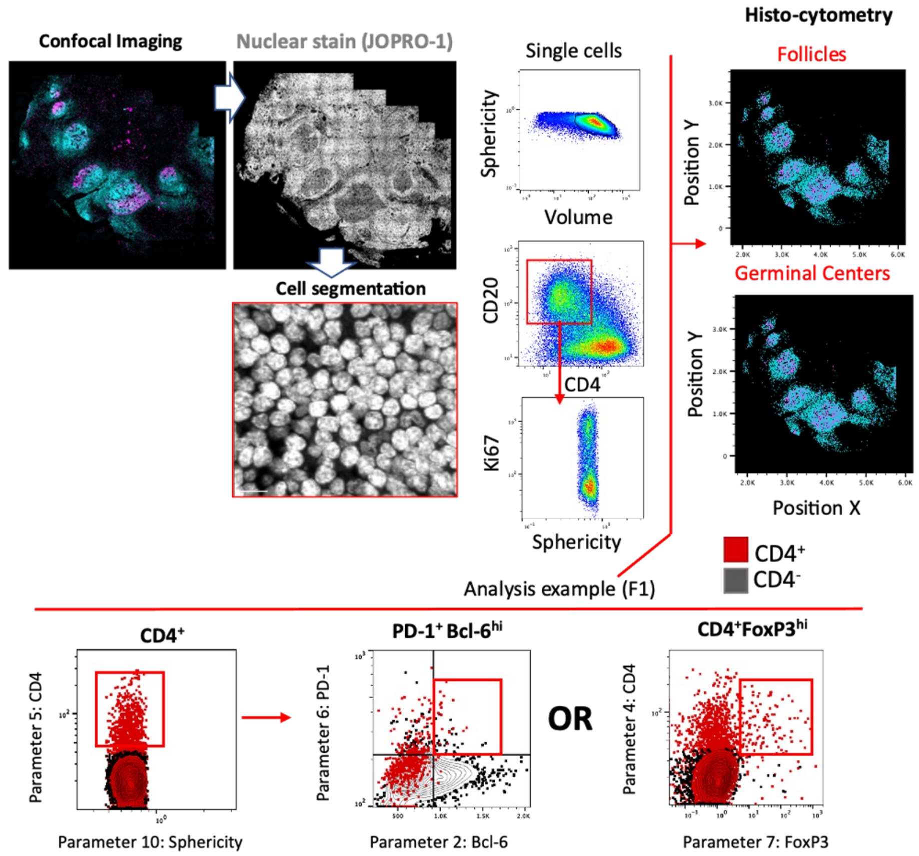

Confocal Imaging
Nuclear stain (JOPRO-1)
Cell segmentation
Single cells
Sphericity
Volume
CD20
CD4
Ki67
Sphericity
Histo-cytometry
Follicles
Germinal Centers
Position Y
Position X
CD4+
CD4-
Analysis example (F1)
CD4+
PD-1+ Bcl-6hi
OR
CD4+FoxP3hi
Parameter 5: CD4
Parameter 10: Sphericity
Parameter 6: PD-1
Parameter 2: Bcl-6
Parameter 4: CD4
Parameter 7: FoxP3

B)

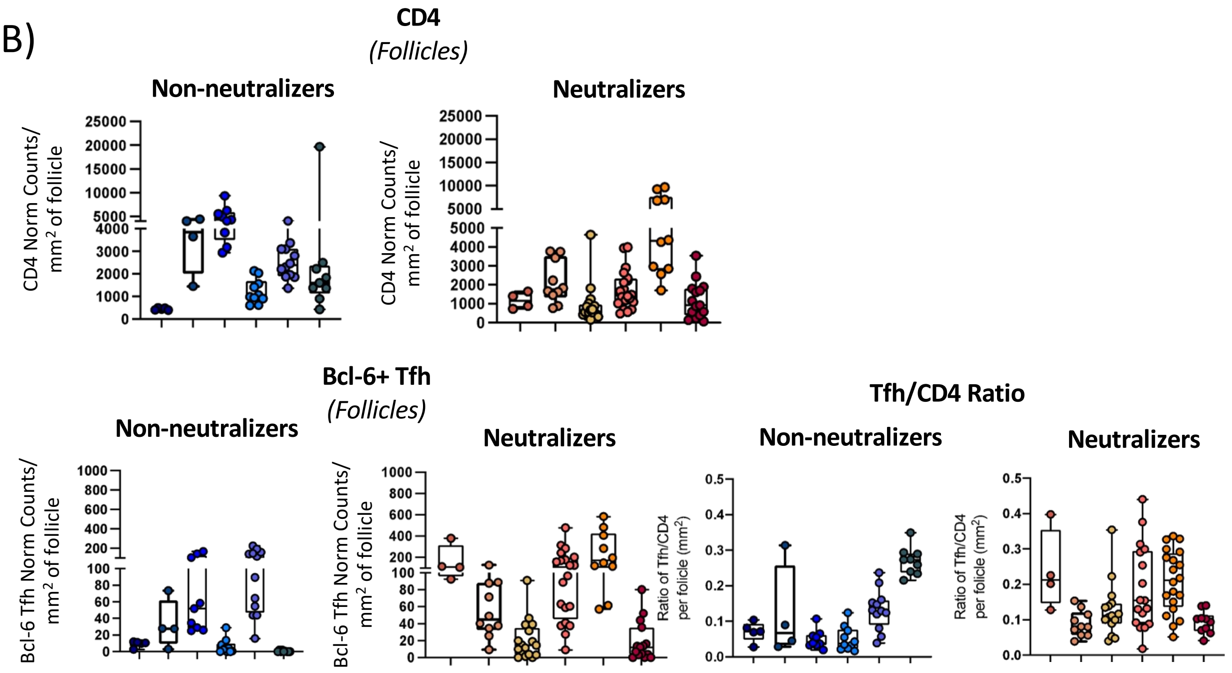

CD4
(Follicles)
Non-neutralizers
Neutralizers
CD4 Norm Counts/ mm2 of follicle
Bcl-6+ Tfh
(Follicles)
Bcl-6 Tfh Norm Counts/ mm2 of follicle
Tfh/CD4 Ratio
Ratio of Tfh/CD4 per folicle (mm2)

**Fig. S3**: ***In situ* characterization of $T_{FH}$ cells**. A) Gating strategy used for subset characterization using Histocytometry. For the analysis, tissues were stained with anti-CD20 and anti-Ki67 for B cell follicle characterization as well as anti-PD1 and anti-Bcl-6 or FoxP3 in combination with CD4 for $T_{FH}$ cells and $T_{FR}$/$T_{REG}$ T cells respectively. The nuclear marker JOPRO-1 was also included to aid with computational segmentation. Confocal images were transformed to FlowJo files as previously described, and CD4, PD-$1^{hi}$Bcl-$6^{hi}$ CD4 (Bcl-6 $T_{FH}$) and FoxP$3^{hi}$ CD4 T cell frequencies within individual B cell follicles were calculated using FlowJo as shown. Original magnification at 40x (NA 1.3). B) Pooled data of normalized CD4 and Bcl-6 $T_{FH}$ cell numbers within the B cell follicles of 6 NN and 6 N LNs. The ratio of $T_{FH}$/CD4 for NNs and Ns is also shown. Each circle represents a single follicle, and each box plot represents tissue from a single donor. Normalization was performed by calculating the number of cells per $mm^2$ of area screened.

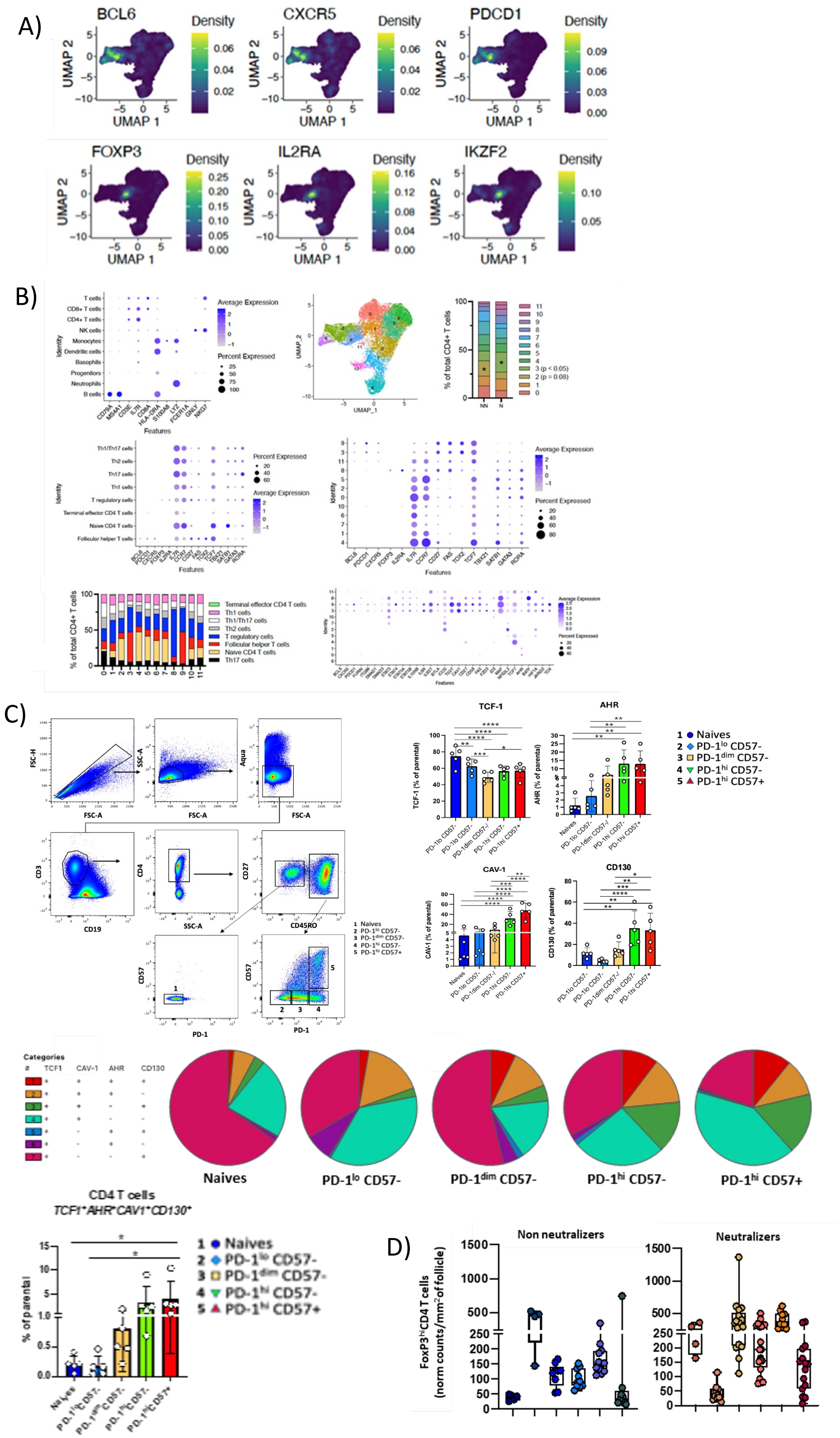
A)
BCL6
CXCR5
PDCD1
FOXP3
IL2RA
IKZF2
Density
UMAP 1
UMAP 2
B)
Average Expression
Percent Expressed
Identity
Features
UMAP_1
UMAP_2
% of total CD4+ T cells
NN
N
3 (p < 0.05)
2 (p = 0.08)
Terminal effector CD4 T cells
Th1 cells
Th1/Th17 cells
Th2 cells
T regulatory cells
Follicular helper T cells
Naive CD4 T cells
Th17 cells
C)
FSC-H
FSC-A
SSC-A
Aqua
CD3
CD19
CD4
CD27
CD45RO
CD57
PD-1
TCF-1
AHR
CAV-1
CD130
1 Naives
2 PD-1lo CD57-
3 PD-1dim CD57-
4 PD-1hi CD57-
5 PD-1hi CD57+
Categories
TCF1
CAV-1
AHR
CD130
Naives
PD-1lo CD57-
PD-1dim CD57-
PD-1hi CD57-
PD-1hi CD57+
CD4 T cells
TCF1+AHR+CAV1+CD130+
% of parental
D)
Non neutralizers
Neutralizers
FoxP3hiCD4 T cells (norm counts/mm2 of follicle)

**Fig. S4**: Identification of CD4 T cell subsets using sc-RNA analysis of LMNCs. A) UMAP plots showing the estimated density of Bcl-6, CXCR5, PDCD1, FOXP3, IL2RA and IKZF2 genes among 11 clusters identified in pooled scRNA-seq analysis. B) Representative marker genes in i) innate immune and adaptive subclusters, ii) CD4 subclusters and iii) 11 subclusters identified in the scRNA-seq analysis. The size of the dots indicates percent cells expressing the gene while the color of the dots indicates the average gene expression level. UMAProjection of CD4 T cells was generated using the Serurat package and SNN clustering was applied to identify 11 clusters within the CD4 T cell compartment. The cluster numbers are shown on the UMAP and the % of each of the 11 identified subclusters within the CD4 T cell compartment is shown using stacked barplots. C) Flow cytometry gating strategy used for the characterization of $CD57^{lo/hi}$ $T_{FH}$ cells and bar charts showing the frequencies of TCF-1, AHR, CAV-1 and CD130 expressing cells within each $T_{FH}$ cell subpopulation as well as combined expression of these four markers and their relative proportion of co-expressing cells (pie charts). Data shown are pooled data from 5 tonsils and each circle represents a different tonsil. D) Cumulative normalized numbers of $FoxP3^{hi}$ CD4 T cells in NNs and Ns as measured by confocal imaging analysis and HistoCytometry. Each circle represents a B cell follicle and follicles from the same donor tissue are grouped together under the same color, with each box plot denoting a different donor.

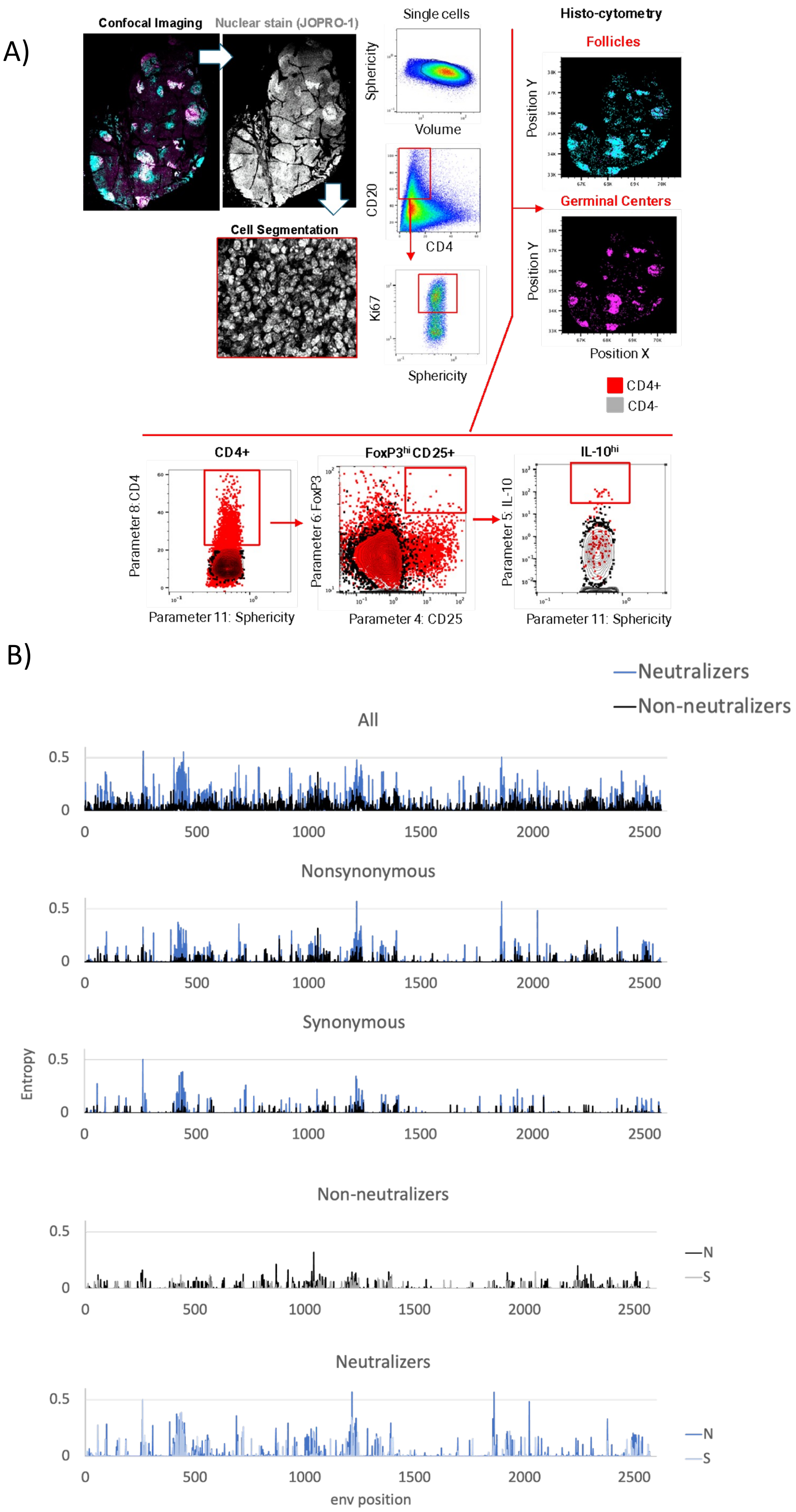
A)
Confocal Imaging
Nuclear stain (JOPRO-1)
Cell Segmentation
Single cells
Sphericity
Volume
CD20
CD4
Ki67
Sphericity
Histo-cytometry
Follicles
Position Y
Germinal Centers
Position Y
Position X
CD4+
CD4-
CD4+
Parameter 8: CD4
Parameter 11: Sphericity
FoxP3hi CD25+
Parameter 6: FoxP3
Parameter 4: CD25
IL-10hi
Parameter 5: IL-10
Parameter 11: Sphericity
B)
Neutralizers
Non-neutralizers
All
Nonsynonymous
Synonymous
Entropy
Non-neutralizers
N
S
Neutralizers
N
S
env position

**Fig. S5:** Regulatory T cell HistoCytometry gating strategy and Entropy-H(x) plots A) Gating strategy used for regulatory T cell characterization by Histocytometry. For the analysis, tissues were stained with anti-CD20 (cyan) and anti-Ki67 (magenta) for B cell follicle characterization as well as anti-CD4, anti-CD25 and anti-FoxP3 in combination with IL-10 for the characterization of corresponding $T_{Fr}$/$T_{REG}$ T cells. The nuclear marker JOPRO-1 was also included to aid with computational segmentation. Confocal images were transformed to FlowJo files as previously described, and $CD25^{hi}FoxP3^{hi}$ IL-$10^{hi}$ CD4 T cell frequencies ($T_{FR}$) within individual B cell follicles were calculated using FlowJo as shown. Original magnification at 40x (NA 1.3)

B) Evolution of HIV Env. Entropy-H(x) plots showing the location of positions with i) all residue mismatches, ii) non-synonymous residue mismatches and iii) synonymous mismatches on alignment to the reference Env HIV sequence in NNs and Ns, and mapping of non-synonymous and synonymous mismatches positions in NNs and NNs.

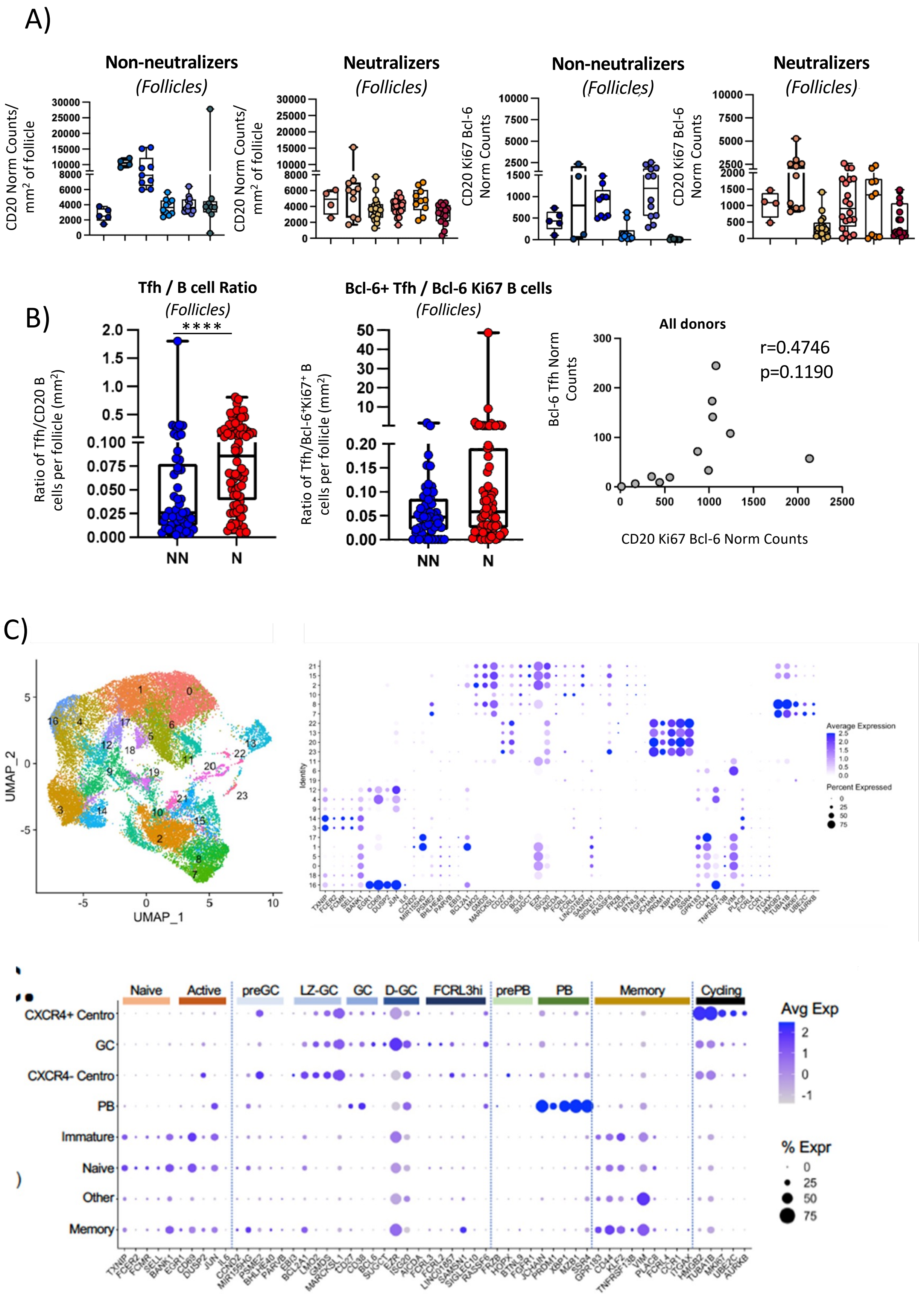
A)
Non-neutralizers
(Follicles)
Neutralizers
(Follicles)
Non-neutralizers
(Follicles)
Neutralizers
(Follicles)
CD20 Norm Counts/ mm² of follicle
CD20 Ki67 Bcl-6 Norm Counts
B)
Tfh / B cell Ratio
(Follicles)
****
Ratio of Tfh/CD20 B cells per follicle (mm²)
NN
N
Bcl-6+ Tfh / Bcl-6 Ki67 B cells
(Follicles)
Ratio of Tfh/Bcl-6⁺Ki67⁺ B cells per follicle (mm²)
NN
N
All donors
r=0.4746
p=0.1190
Bcl-6 Tfh Norm Counts
CD20 Ki67 Bcl-6 Norm Counts
C)
UMAP_2
UMAP_1
Average Expression
Percent Expressed
Identity
Naive
Active
preGC
LZ-GC
GC
D-GC
FCRL3hi
prePB
PB
Memory
Cycling
CXCR4+ Centro
GC
CXCR4- Centro
PB
Immature
Naive
Other
Memory
Avg Exp
% Expr

**Fig. S6**: Identification of B cell clusters using sc-RNA analysis of LMNCs. A) Box plots showing the normalized numbers of $CD20^{hi}$ and $CD20^{hi}Ki67^{hi}Bcl\text{-}6^{hi}$ B cells in NNs and Ns as measured by Histocytometry. Each donor is represented by a different color. Circles represent single follicles and follicles from the same tissue share the same color. B) Pooled analysis of B cell follicle $T_{FH}$ cell/CD20 B cell ratios (left: ****$p<0.0001$) and $T_{FH}$ cells / $Bcl\text{-}6^{hi}Ki67^{hi}$ B cells ($p=0.0568$) in NNs and Ns as measured by confocal imaging and Histocytometry. Each circle represents a single B cell follicle and follicles from the same group are shown together (NNs: blue, Ns: red). The correlation between $Bcl\text{-}6^{hi}$ $T_{FH}$ and $Bcl\text{-}6^{hi}Ki67^{hi}CD20^{hi}$ B cells numbers is shown too. Group differences between NNs and Ns were determined by One-Way ANOVA. Probability values $p<0.05$ were considered significant. C) UMAP projection of 23 B-cell subclusters identified through pooled scRNA-seq analysis (left) and their representative marker genes (right). The immune phenotypes corresponding to these 23 B cell subclusters are also described. The size of the dots indicates percent cells expressing the gene while the color of the dots indicates the average gene expression level.

**Table S1:** Clinical Information of study participants

| ID | Age | Gender | $VL_{log}$ | %CD4 | [CD4 count] | Treatment | Tissue Type |
|---|---|---|---|---|---|---|---|
| Non neutralizers | | | | | | | |
| HIV #1 | 24 | M | 5.51 | 716 | 24 | ART-naive | Cervical LN |
| HIV #2 | 30 | F | 1.6 | 438 | 37 | ART-naive | Cervical LN |
| HIV #3 | 34 | M | 5.33 | 425 | 17 | ART-naive | Cervical LN |
| HIV #4 | 18 | M | 1.6 | 504 | 32 | ART-naive | Cervical LN |
| HIV #5 | 31 | M | 6.3 | 424 | 20 | ART-naive | Cervical LN |
| HIV #6 | 29 | M | 4.46 | 1288 | 37 | ART-naive | Cervical LN |
| HIV #7 | 23 | M | 5.06 | 319 | 25 | ART-naive | Cervical LN |
| HIV #8 | 23 | M | 6.08 | 315 | 9 | ART-naive | Cervical LN |
| HIV #9 | 28 | M | 4.65 | 382 | 21 | ART-naive | Cervical LN |
| HIV #10 | 40 | M | 5.61 | 153 | 11 | ART-naive | Cervical LN |
| HIV #11 | 20 | F | 3.85 | 504 | 27 | ART-naive | Cervical LN |
| HIV #12 | 22 | M | 7 | 381 | 27 | ART-naive | Cervical LN |
| HIV #13 | 19 | M | 5.13 | 212 | 17 | ART-naive | Cervical LN |
| HIV #14 | 28 | M | 4.59 | 394 | 24 | ART-naive | Cervical LN |
| HIV #15 | 24 | M | 5.79 | 290 | 13 | ART-naive | Inguinal LN |
| HIV #16 | 31 | M | 4.79 | 406 | 25 | ART-naive | Inguinal LN |
| Neutralizers | | | | | | | |
| HIV #17 | 24 | M | 5.25 | 4 | 1 | ART-naive | Cervical LN |
| HIV #18 | 20 | M | 5.76 | 175 | 8 | ART-naive | Cervical LN |
| HIV #19 | 40 | F | 5.05 | 279 | 18 | ART-naive | Cervical LN |
| HIV #20 | 61 | M | 5.65 | 321 | 7 | ART-naive | Cervical LN |
| HIV #21 | 48 | M | 5.13 | 415 | 10 | ART-naive | Cervical LN |
| HIV #22 | 31 | M | 4.58 | 591 | 15 | ART-naive | Inguinal LN |
| HIV #23 | 38 | M | 4.85 | 387 | 20 | ART-naive | Cervical LN |
| HIV #24 | 31 | M | 4.24 | 274 | 22 | ART-naive | Cervical LN |
| HIV #25 | 39 | M | 4.96 | 179 | 9 | ART-naive | Cervical LN |
| HIV #26 | 38 | M | 4.76 | 40 | 1 | ART-naive | Cervical LN |
| HIV #27 | 40 | M | 5.5 | 64 | 7 | ART-naive | Cervical LN |
| HIV #28 | 22 | M | 4.42 | 135 | 12 | ART-naive | Cervical LN |
| HIV #29 | 23 | M | 5.13 | 285 | 21 | ART-naive | Cervical LN |
| HIV #30 | 27 | M | 4.15 | 671 | 21 | ART-naive | Inguinal LN |
| HIV #31 | 32 | M | 3.65 | 548 | 18 | ART-naive | Inguinal LN |
| HIV #32 | 31 | M | 4.64 | 228 | 12 | ART-naive | Inguinal LN |
| HIV #33 | 29 | M | 5.27 | 348 | 12 | ART-naive | Cervical LN |
| HIV negative | | | | | | | |
| #34_1E | 77 | M | n/a | | | n/a | Axillary LN |
| #35_1F | 44 | M | n/a | | | n/a | Cervical LN |
| #36_1G | 52 | M | n/a | | | n/a | Inguinal LN |
| #37_1H | 28 | F | n/a | | | n/a | Inguinal LN |
| #38_1U | 78 | M | n/a | | | n/a | Inguinal LN |
| #39_1J | 22 | F | n/a | | | n/a | Cervical LN |

**Table S2:** Study Assays

| ID | Tissue Type | Plasma Ab Fingerprinting | Flow | scRNA | mIF_Tfh/B | mIF_Treg | mIF_RNA | HIVseq |
|---|---|---|---|---|---|---|---|---|
| **Non neutralizers** | | | | | | | | |
| HIV #1 | Cervical LN | | | | ● | ● | ● | ● |
| HIV #2 | Cervical LN | | ● | | | | | ● |
| HIV #3 | Cervical LN | | | | ● | ● | ● | |
| HIV #4 | Cervical LN | | | | | | | |
| HIV #5 | Cervical LN | | ● | ● | | | | |
| HIV #6 | Cervical LN | | ● | | ● | ● | ● | ● |
| HIV #7 | Cervical LN | | ● | | | | | |
| HIV #8 | Cervical LN | | ● | | | | | |
| HIV #9 | Cervical LN | | | | ● | ● | ● | ● |
| HIV #10 | Cervical LN | | ● | | | | | ● |
| HIV #11 | Cervical LN | | ● | | | | | ● |
| HIV #12 | Cervical LN | | ● | | ● | ● | ● | |
| HIV #13 | Cervical LN | | ● | ● | ● | ● | ● | ● |
| HIV #14 | Cervical LN | | ● | | | | | ● |
| HIV #15 | Inguinal LN | | | ● | | | | ● |
| HIV #16 | Inguinal LN | | | ● | | | | ● |
| **Neutralizers** | | | | | | | | |
| HIV #17 | Cervical LN | ● | ● | | | | | ● |
| HIV #18 | Cervical LN | ● | ● | ● | | | | ● |
| HIV #19 | Cervical LN | ● | ● | ● | | | | ● |
| HIV #20 | Cervical LN | ● | ● | | ● | ● | ● | ● |
| HIV #21 | Cervical LN | ● | ● | | ● | ● | ● | ● |
| HIV #22 | Inguinal LN | ● | | | ● | ● | ● | |
| HIV #23 | Cervical LN | ● | | | ● | | ● | |
| HIV #24 | Cervical LN | ● | | | ● | ● | ● | |
| HIV #25 | Cervical LN | ● | | | | | | |
| HIV #26 | Cervical LN | ● | | | | | | |
| HIV #27 | Cervical LN | ● | | | | | | |
| HIV #28 | Cervical LN | ● | | | | | | |
| HIV #29 | Cervical LN | ● | | | | | | |
| HIV #30 | Inguinal LN | ● | ● | ● | ● | ● | ● | ● |
| HIV #31 | Inguinal LN | ● | | | | | | |
| HIV #32 | Inguinal LN | ● | ● | ● | | | | |
| HIV #33 | Cervical LN | ● | | | | | | ● |
| **HIV negative** | | | | | | | | |
| #34_1E | Axillary LN | | | | ● | | | |
| #35_1F | Cervical LN | | | | ● | | | |
| #36_1G | Inguinal LN | | | | ● | | | |
| #37_1H | Inguinal LN | | | | ● | | | |
| #38_1U | Inguinal LN | | | | ● | | | |
| #39_1J | Cervical LN | | | | ● | | | |

**Table S3:** Flow Cytometry Antibodies

| Target | Fluorophore | Clone | Manufacturer |
|---|---|---|---|
| **Tfh Panel** | | | |
| CD4 | BUV805 | SK3 | BD Biosciences |
| CD3 | H7APC | SK7 | BD Biosciences |
| CD27 | A700 | M-T271 | BD Biosciences |
| CD45RO | BV786 | UCHL1 | BD Biosciences |
| TIM-3 (CD366) | BUV661 | 7D3 | BD Biosciences |
| CD8 | BV570 | RPA-T8 | Biolegend |
| CD19 | Cy55PE | SJ25C1 | Invitrogen |
| PD-1 (CD279) | BUV496 | EH12.1 | BD Biosciences |
| CXCR5 (CD185) | Cy7PE | MU5UBEE | Invitrogen |
| TIGIT | BB660 | MB5A43 | BD Biosciences |
| CD95 | BV480 | DX2 | BD Biosciences |
| CXCR3 (CD183) | BV650 | G025H7 | Biolegend |
| ICOS (CD278) | BV421 | DX29 | BD Biosciences |
| PDL-1 (CD274) | BB515 | MIH1 | BD Biosciences |
| CD226 | BB700 | DX11 | BD Biosciences |
| CD57 | BB790 | NK-1 | BD Biosciences |
| OX40 (CD134) | BV711 | L106 | BD Horizon |
| CTLA-4 (CD152) | PE | L3D10 | Biolegend |
| OX40L (CD252) | BUV563 | iK-1 | BD Biosciences |
| **Transcription Factor Panel** | | | |
| CD4 | BV650 | SK3 | BD Horizon |
| CD3 | H7APC | SK7 | BD Biosciences |
| CD45RO | BV786 | UCHL1 | BD Biosciences |
| CD27 | BV605 | O323 | BD Biosciences |
| PD-1 (CD279) | BV711 | EH12.2H7 | Biolegend |
| CD57 | BB790 | NK-1 | BD Biosciences |
| CD19 | ECD | J3-119 | Beckman-Coulter |
| AHR | AF488 | polyclonal | R&D systems |
| CAV-1 | PE | Polyclonal | Abcam |
| TCF1 | AF647 | 7F11A10 | Biolegend |
| CD130 | BV421 | AM64 | BD Biosciences |

**Table S4:** Multiparameter Imaging Antibodies

| Target | Fluorophore | Species | Isotype | Clone | Manufacturer # Catalog No |
|---|---|---|---|---|---|
| **Tfh Panel** | | | | | |
| Bcl-6 | Unconjugated | Mouse | IgG1 | PG-B6p | DAKO, #M7211 |
| PD-1 | Alexa Fluor 488 | Mouse | IgG | polyclonal | R&D systems, #FAB7115G |
| Ki67 | Brilliant Violet 421 | Mouse | IgG1 | B56 | BD Horizon, #562899 |
| CD20 | eFluor 615 | Mouse | IgG2a | L26 | eBioscience, #42-0202-82 |
| FoxP3 | Alexa Fluor 647 | Mouse | IgG1 | 206D | Biolegend, #320114 |
| CD4 | Alexa Fluor 700 | Goat | IgG | polyclonal | R&D systems, # FAB8165N |
| **RNAscope panel** | | | | | |
| PD-1 | Unconjugated | Rabbit | IgG | EPR4877 | Abcam, #ab137132 |
| FDC | Unconjugated | Mouse | IgM | F3803 | SIGMA, #F3803 |
| CD3 | Unconjugated | Mouse | IgG1 | F7.2.38 | DAKO, #M7254 |
| CD20 | eFluor 615 | Mouse | IgG2a | L26 | eBioscience, #42-0202-82 |
| CD4 | Alexa Fluor 700 | Goat | IgG | polyclonal | R&D systems, # FAB8165N |
| **Transcription Factor Panel** | | | | | |
| CD25 | unconjugated | Mouse | IgG2b | 4C9 | BioSB, # BSB 6320 |
| IL-10 | Alexa Fluor 546 | Mouse | IgG2b | E-10 | Santa Cruz, #sc-8438 |
| Ki67 | Brilliant Violet 480 | Mouse | IgG1 | B56 | BD Horizon, # 566172 |
| CD20 | eFluor 615 | Mouse | IgG2a | L26 | eBioscience, #42-0202-82 |
| FoxP3 | Alexa Fluor 647 | Mouse | IgG1 | 206D | Biolegend, #320114 |
| CD4 | Alexa Fluor 700 | Goat | IgG | polyclonal | R&D systems, # FAB8165N |
| **Secondary antibodies (all panels)** | | | | | |
| Goat | Alexa Fluor 546 | Mouse | IgG1 | n/a | Life Technologies, # A21121 |
| Donkey | Brilliant Violet 421 | Rabbit | IgG | Poly4046 | Biolegend, #406410 |
| Goat | Alexa Fluor 546 | Mouse | IgM | n/a | Life Technologies, #A21123 |
| Goat | Alexa Fluor 488 | Mouse | IgG2b | n/a | Life Technologies, #A21141 |
| Goat | Alexa Fluor 488 | Mouse | IgG1 | n/a | Life Technologies, #A21121 |
